# Nitric oxide prevents aortic valve calcification by S-nitrosylation of USP9X to activate NOTCH signaling

**DOI:** 10.1101/2020.08.15.252171

**Authors:** Uddalak Majumdar, Sathiyanarayanan Manivannan, Madhumita Basu, Yukie Ueyama, Mark C. Blaser, Emily Cameron, Michael R. McDermott, Joy Lincoln, Susan E. Cole, Stephen Wood, Elena Aikawa, Brenda Lilly, Vidu Garg

**Affiliations:** Center for Cardiovascular Research, Nationwide Children’s Hospital, Columbus, Ohio, USA; The Heart Center, Nationwide Children’s Hospital, Columbus, Ohio, USA; Department of Pediatrics, The Ohio State University, Columbus, Ohio, USA; Division of Cardiovascular Medicine, Department of Medicine, Center for Interdisciplinary Cardiovascular Sciences Brigham and Women’s Hospital, Harvard Medical School, Boston, MA, USA; Division of Cardiovascular Medicine, Department of Medicine, Center of Excellence in Cardiovascular Biology, Brigham and Women’s Hospital, Harvard Medical School, Boston, MA, USA; Department of Pediatrics, Medical College of Wisconsin, Division of Pediatric Cardiology, Children’s Wisconsin, Milwaukee, WI, USA; Herma Heart Institute, Division of Pediatric Cardiology, Children’s Wisconsin, Milwaukee, WI, USA; Griffith Institute for Drug Discovery, Griffith University, Brisbane, Queensland, Australia; Department of Molecular Genetics, The Ohio State University, Columbus, Ohio, USA

**Keywords:** Calcific aortic valve disease, NOTCH1, Nitric oxide, S-nitrosylation, USP9X

## Abstract

Calcific aortic valve disease (CAVD) is an increasingly prevalent condition and endothelial dysfunction is implicated in its etiology. We previously identified nitric oxide (NO) as a calcification inhibitor by its activation of *NOTCH1*, which is genetically linked to human CAVD. Here, we show that NO rescues calcification by a S-nitrosylation-mediated mechanism in porcine aortic valve interstitial cells (pAVICs) and single cell RNA-seq demonstrated regulation of NOTCH pathway by NO. A unbiased proteomic approach to identify S-nitrosylated proteins in valve cells found enrichment of the ubiquitin proteasome pathway and implicated S-nitrosylation of USP9X in NOTCH regulation during calcification. Furthermore, S-nitrosylated USP9X was shown to deubiquitinate and stabilize MIB1 for NOTCH1 activation. Consistent with this, genetic deletion of *Usp9x* in mice demonstrated aortic valve disease and human calcified aortic valves displayed reduced S-nitrosylation of USP9X. These results demonstrate a novel mechanism by which S-nitrosylation dependent regulation of ubiquitin-associated pathway prevents CAVD.

---

Global prevalence of calcific aortic valve disease (CAVD) is 12.6 million per year and affects ∼2% of individuals over 70 years of age^1^. Inadequate cardiac output due to left ventricular outflow obstruction caused by the stenotic aortic valve results in decreased exercise capacity, progressive left ventricular hypertrophy and ultimately heart failure and death^2^. CAVD is an active and progressive condition and is the main contributor to the development of aortic valve stenosis in adults^2^. Due to limited molecular understanding of disease mechanisms, no pharmacological treatment is available to effectively prevent the onset or progression of CAVD and aortic valve replacement remains the only treatment option^3,4^.

A healthy aortic valve (AoV) is able to open fully during systole and close during diastole to create unidirectional blood flow. In CAVD, the valve leaflets cannot open to their full extent due to increased stiffness. The diseased AoV is characterized by thick, fibrotic leaflets, often covered with calcific nodules on the aortic surface^2^. Each valve leaflet is composed of three stratified layers of specialized extracellular matrix (ECM), which provide biomechanical strength in order to withstand a constant oscillating hemodynamic environment. The ECM is interspersed with valve interstitial cells (VICs) and encapsulated by a monolayer of valve endothelial cells (VECs). There is bidirectional communication between VECs and VICs, which regulates the behavior of each cell type^5,6^. Turnover of the valve ECM is regulated by VICs, which are largely quiescent and fibroblast-like in healthy valves^7,8^. CAVD is proposed to be driven by a phenotypic switch of VICs from fibroblast-like to myofibroblast and osteoblast-like cells that deposit bone-like matrix restricting the movement of valve cusps. Valvular heart disease is often triggered by injury and/or impaired VEC-VIC signaling^6,9^.

We previously demonstrated that VEC-derived nitric oxide (NO), a small lipophilic second messenger molecule, plays an important role in AoV development and disease including VIC calcification^10^. *In vivo* and *in vitro* studies demonstrated that NO can activate the NOTCH1 signaling pathway to inhibit calcification, and mutations in *NOTCH1* were associated with bicuspid aortic valve and CAVD in humans^10,11^. Our laboratory established a genetic interaction between endothelial NO synthase and *Notch1* in mice that was important for CAVD pathogenesis^10^. However, the mechanistic basis of NO-mediated NOTCH1 activation and protection from CAVD was not defined. NO exerts most of its cellular influence either by sGC/cGMP pathway activation or S-nitrosylation, a reversible post-translational modification of Cys-thiol group^12^. S-nitrosylation regulates a wide range of critical cellular functions including metabolism, membrane trafficking, enzyme activity and stability through both allosteric and active-site modification^13^. In addition, S-nitrosylation also regulates other post-translational modifications such as phosphorylation and ubiquitination, which in turn influence protein function and degradation that are important for cardiovascular and various other diseases^13,14^. The ubiquitin-proteasome pathway has been implicated in almost every aspect of cellular function and signaling in eukaryotes^15^. Not surprisingly, the ubiquitin-proteasome pathway also regulates aspects of the NOTCH signaling pathway^16,17^, which is important for AoV development and progression of disease^11^. However, the correlation between these two pathways in the development of valve disease is not clear.

Here, we demonstrated for the first time how S-nitrosylation protects the valve leaflet from calcification by inhibiting myofibroblastic activation of aortic VICs (AVICs). Mass spectrometry analysis of S-nitrosylated proteins in AVICs revealed involvement of the ubiquitin proteasome pathway in the calcific process. We identified S-nitrosylation mediated activation of a novel deubiquitinase, USP9X, a NOTCH1 signaling regulator in AVICs. We further showed that S-nitrosylation of USP9X stabilizes MIB1, which activates NOTCH1 signaling in neighboring cells to prevent calcification. Our data also revealed the involvement of USP9X in CAVD using a murine model harboring a conditional deletion of *Usp9x* in endothelial and endothelial-derived cells. In humans, we also observed reduced S-nitrosylation of USP9X in calcified AoVs with an associated reduction in MIB1 and activated-NOTCH1 expression, consistent with the proposed mechanism. Overall, we demonstrated a novel mechanism that prevents CAVD mediated by S-nitrosylation and the ubiquitin proteasome pathway.

## Results

### Nitric oxide inhibits calcification in AVICs via S-nitrosylation

We and others previously reported that nitric oxide (NO) donor inhibits spontaneous calcification of cultured porcine aortic valve interstitial cells (pAVICs) *in vitro*^10,18^. Additionally, our laboratory found that the presence of NO donor was associated with the activation of Notch1 signaling, but the underlying mechanism was not known^10^. NO functions either by activating the sGC/cGMP pathway or by modulating redox-dependent, post-translational modification, such as S-nitrosylation. In order to define the NO-mediated mechanism involved in the inhibition of calcification, we specifically activated either the sGC/cGMP pathway, by adding cGMP analog 8-Br-PET-cGMP, or S-nitrosylation, with the addition of S-nitrosoglutathione (GSNO) (Figure 1). While inhibition of spontaneous calcification of pAVICs was noted in the presence of NO donor, calcific nodules were continued to be observed after activation of the sGC/cGMP pathway suggesting that NO-mediated inhibition of calcification is independent of this pathway (Figure 1 A-C, E). Similar to the NO donor treatment, calcification was blocked in the presence of a S-nitrosylating agent (GSNO), consistent with the involvement of S-nitrosylation in the inhibition of calcification (Figure 1 D, E). RUNX2, a marker of osteoblast differentiation, was downregulated in the presence of NO donor and GSNO, but unaltered after activation of the sGC/cGMP pathway with 8-Br-PET-cGMP (Figure 1 F-J; Supplemental Figure 1 A, B). We confirmed the activation of sGC/cGMP pathway by measuring phosphorylated vasodilator-stimulated phosphoprotein (pVASP), a known target of sGC/cGMP pathway^19^. pVASP was increased in the presence of 8-Br-PET-cGMP, but its expression was unchanged in the presence of NO donor and GSNO, indicating no activation of sGC/cGMP pathway (Supplemental Figure 1).

**Figure 1.**
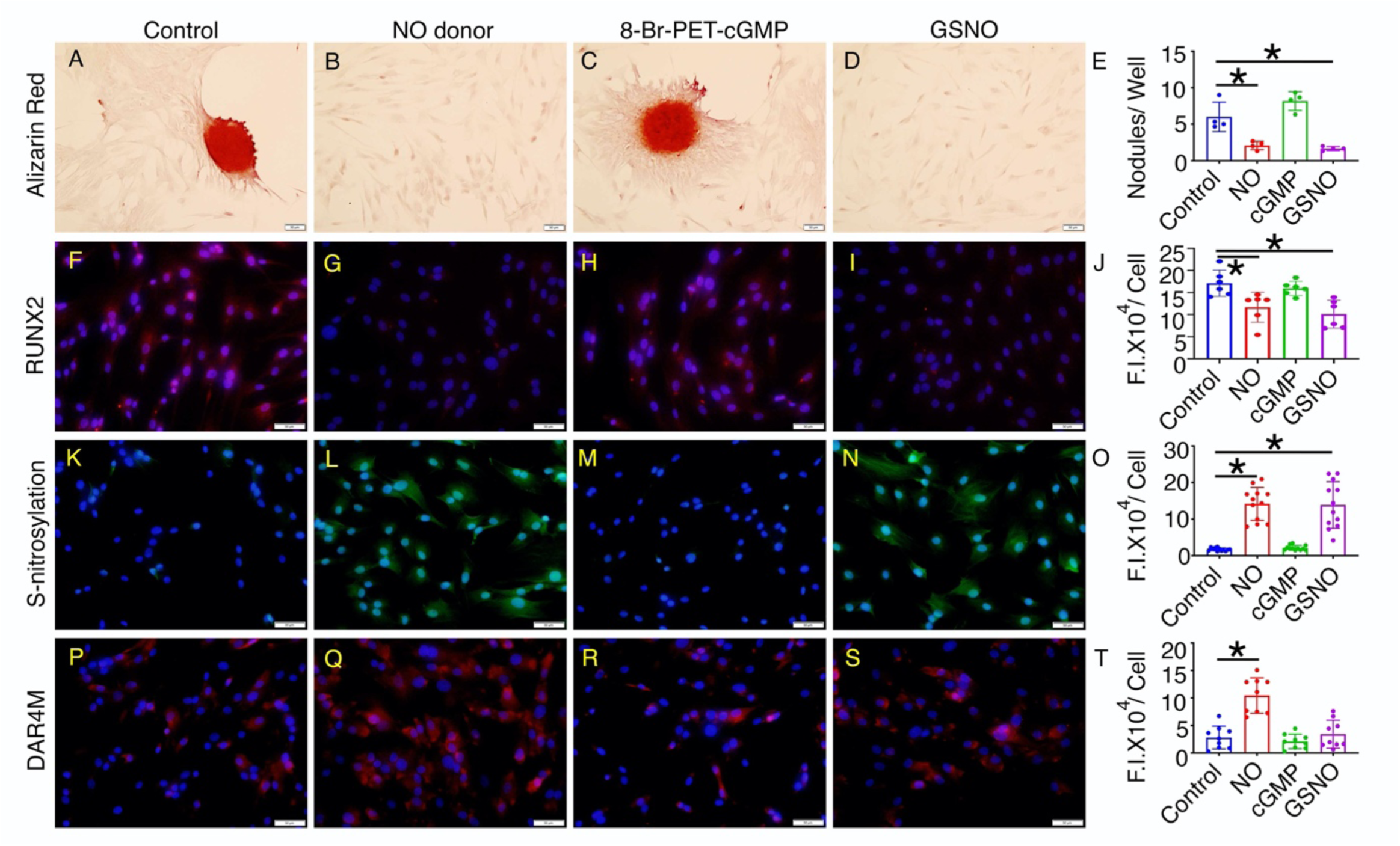
Calcification of porcine aortic valve interstitial cells (AVICs) is attenuated with addition of detaNONOate (NO donor) or GSNO (S-nitrosylating agent). (A-E) Alizarin Red staining and nodule formation of porcine AVICs cultured for 5 days under control conditions or with addition of NO donor, 8-Br-PET-cGMP (cGMP analog) or GSNO. (F-I) RUNX2 (red) expression with nuclear DAPI staining (blue) is shown in four conditions and quantified in (J). (K-N) represent total cellular S-nitrosylated proteins (green) counterstained with DAPI and is quantified in (O). (P-S) DAR4M staining (red) which measures NO is shown with DAPI (blue) counterstaining and is quantified in (T). Scale bar: 50 μm. * represent *P* value ≤ 0.05 (2-tailed). For all experiments, Mann-Whitney test was performed. F.I.: Fluorescence Intensity.

Furthermore, we found evidence of increased S-nitrosylation of proteins in pAVICs treated with NO donor and GSNO compared to untreated cells or those treated with 8-Br-PET-cGMP, which suggests S-nitrosylation is important for NO-mediated suppression of calcification in pAVICs (Figure 1 K-O). We also assessed the amount of NO in pAVICs by DAR4M staining in the unlikely event that GSNO was degraded into glutathione (GSH) and NO to activate sGC/cGMP pathway. NO was significantly increased after addition of NO donor and was not significantly altered after treatment with either GSNO or 8-Br-PET-cGMP (Figure 1 P-T).

### Nitric oxide inhibits pAVIC-activation and regulates NOTCH1 target genes

We next evaluated the transcriptional changes that occurred in cultured pAVICs with the addition of NO donor using single cell RNA-seq (scRNAseq). The gene expression of pAVICs cultured in the presence of NO donor for five days were compared to untreated pAVICs. Unbiased clustering of the gene expression from both of these cell populations using UMAP demonstrated two distinct populations and suggested that the effect of NO on pAVICs is uniform and not specific to a subpopulation of cells (Figure 2A).

**Figure 2.**
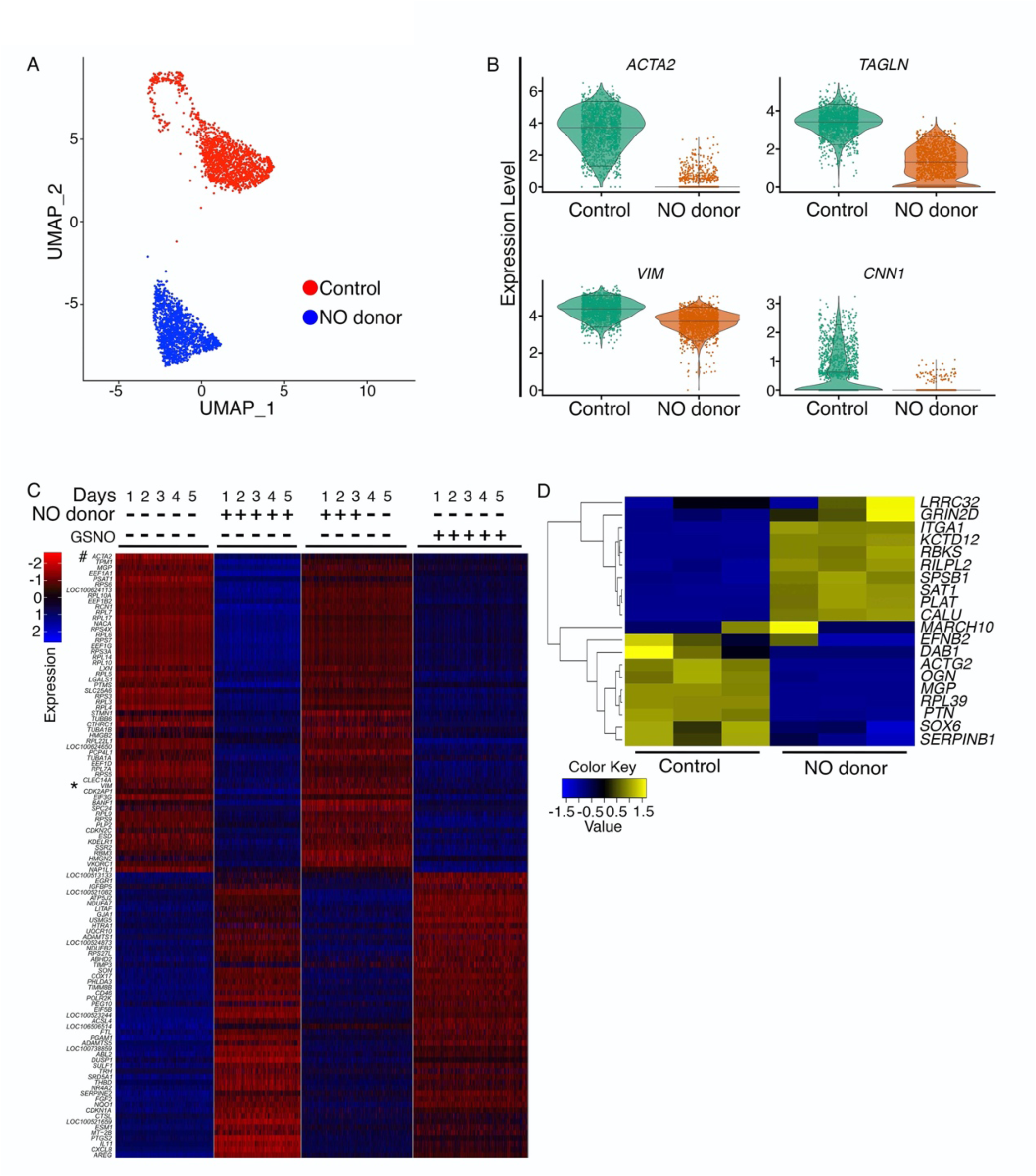
Gene expression analysis of porcine aortic valve interstitial cells (AVICs) exposed to NO donor or GSNO by single cell RNA sequencing (scRNA-seq). (A) UMAP plot of scRNA-seq expression data shows two distinct populations of porcine AVICs cultured in the presence (blue) or absence (red) of NO donor. (B) Violin plots show downregulation of *ACTA2, TAGLN, VIM* and *CNN1* with NO donor. Each dot in violin plot represents gene expression in individual cells. Horizontal lines represent 0.05, 0.5 and 0.95 quantiles of gene expression. (C) Heatmap represents Z-scored expression of top 100 differentially expressed genes in individual cells from the four culture conditions on day five. # and * indicates *ACTA2* and *VIM*, respectively. (D) Heatmap demonstrating Z-scored, log2-transformed expression (CPM) of NOTCH1 target genes in porcine AVICs in presence and absence of NO donor.

To confirm that pAVICs were representative of VICs *in vivo*, we compared our *in vitro* scRNAseq data from pAVICs to published scRNAseq data of one-month old (P30) mouse AoV cells^20^. We observed significant correlation between the expression of mouse AVIC marker genes, *Col1a1, Col3a1* and *Serpinf1*, with pAVICs and the weak expression of endothelial (*Cdh5, Nos3*, and *Emcn*), melanocyte (*Dct, Mitf, Tyrp1*) or immune cell (*Tyrobp, Ptprc, Cx3cr1*) genes in pAVICs (Supplemental Figure 2A). This observation indicates that our pAVICs culture is homogeneous and does not contain a significant percentage of other cell types found within the adult aortic valve.

Differential expression analysis of scRNAseq data of pAVICs revealed *ACTA2* (α-SMA) expression is strongly suppressed in pAVICs cultured in presence of NO donor (Figure 2B, Supplemental Figure 2B). As *ACTA2* is a characteristic marker of myofibroblast differentiation, this decrease demonstrates that NO suppresses conversion of AVICs into myofibroblast-like cells. In addition, other myofibroblast markers, such as *TAGLN* (Transgelin), *VIM* (Vimentin) and *CNN1* (Calponin)^21^, were also reduced, further supporting that NO inhibits myofibroblast activation of pAVICs (Figure 2B, Supplemental Figure 2B). To test whether this NO mediated inhibition of myofibroblast activation is transient, pAVICs were maintained in presence of NO donor for first three days followed by culturing cells without NO donor for additional two days and performed scRNAseq. Interestingly, RNA expression profile after NO donor withdrawal is similar to that of the pAVICs cultured in absence of NO donor (Figure 2C). This suggests that effect of NO on gene expression is not permanent, and cellular markers of a myofibroblast-like state, including *ACTA2* and *VIM* rapidly reappear in these cells (Figure 2C, Online data I). To examine the gene expression after S-nitrosylation, we cultured pAVICs for five days in presence of GSNO followed by scRNAseq. We observed gene expression profile after GSNO exposure is similar to NO donor exposure implying that NO-mediated changes to gene expression in pAVICs are in large part due to S-nitrosylation (Figure 2C). Consistent with this mechanism, expression of *GUCY1A3* and *GUCY1B3*, which encodes soluble guanylate cyclase (sGC) were not significantly affected after addition of NO donor (Supplemental Figure 2B). This observation further indicates that sGC/cGMP pathway is not involved in NO-mediated inhibition of myofibroblast activation and calcification of pAVICs, and the effect of NO on calcification of pAVICs is due to S-nitrosylation.

Our previous studies have shown that NOTCH signaling is increased by NO treatment, as demonstrated by increased nuclear localization of the NOTCH1 intracellular domain (NICD)^10^. Examination of scRNA-seq data demonstrated that *NOTCH1* mRNA expression was not increased after NO treatment (Supplemental Figure 2B). This prompted us to examine if NOTCH1 signaling targets were affected by NO treatment. We compared pAVICs scRNAseq data with published NOTCH1 target genes that were identified by combining ChIP-seq and RNAseq datasets from *NOTCH1*-siRNA treated human AVECs subjected to shear stress^22^. We observed changes of NOTCH1 target genes in pAVICs after NO exposure similar to that observed in human AVECs in response to shear stress (Figure 2D; Supplemental Figure 2C). This similarity of gene expression confirms that NOTCH signaling is impacted by NO treatment in pAVICs. The absence of *NOTCH1* mRNA changes at the transcriptional level, but alterations in NOTCH1 signaling targets supported a post-translational mechanism for the regulation of NOTCH1 signaling by NO. Based upon the transcriptomic changes observed with GSNO treatment, S-nitrosylation was the most likely post-translational mechanism for this regulation.

### USP9X, an activator of NOTCH1 signaling, is S-nitrosylated in pAVICs

In order to identify proteins which were S-nitrosylated after exposure to NO donor in AVICs, we utilized a modified biotin switch technique (BST) followed by mass spectrometry (LC/MS-MS). Rat AVICs were used for the LC/MS-MS identification of S-nitrosylated proteins as the porcine protein sequences are not well annotated. Similar to pAVICs, rat AVICs undergo calcification and the addition of NO donor inhibited their osteogenic differentiation, as demonstrated by RUNX2 expression (Supplemental Figure 3A). Modified BST followed by LC/MS-MS identified total 579 S-nitrosylated proteins in rat AVICs (untreated control and NO donor treated samples) and among them, 217 proteins were over-represented after NO treatment (Supplemental figure 3B, C; Online data II-IX). Enrichment analysis of S-nitrosylated proteins utilizing WebGestalt 2019 (WEB-based Gene SeT AnaLysis Toolkit)^23^ revealed that ubiquitin specific processing proteases was the most enriched pathway after NO treatment. Additionally, the deubiquitination pathway, representing 12 similar proteins as identified for ubiquitin specific processing proteases, was also over-represented (Figure 3A).

**Figure 3.**
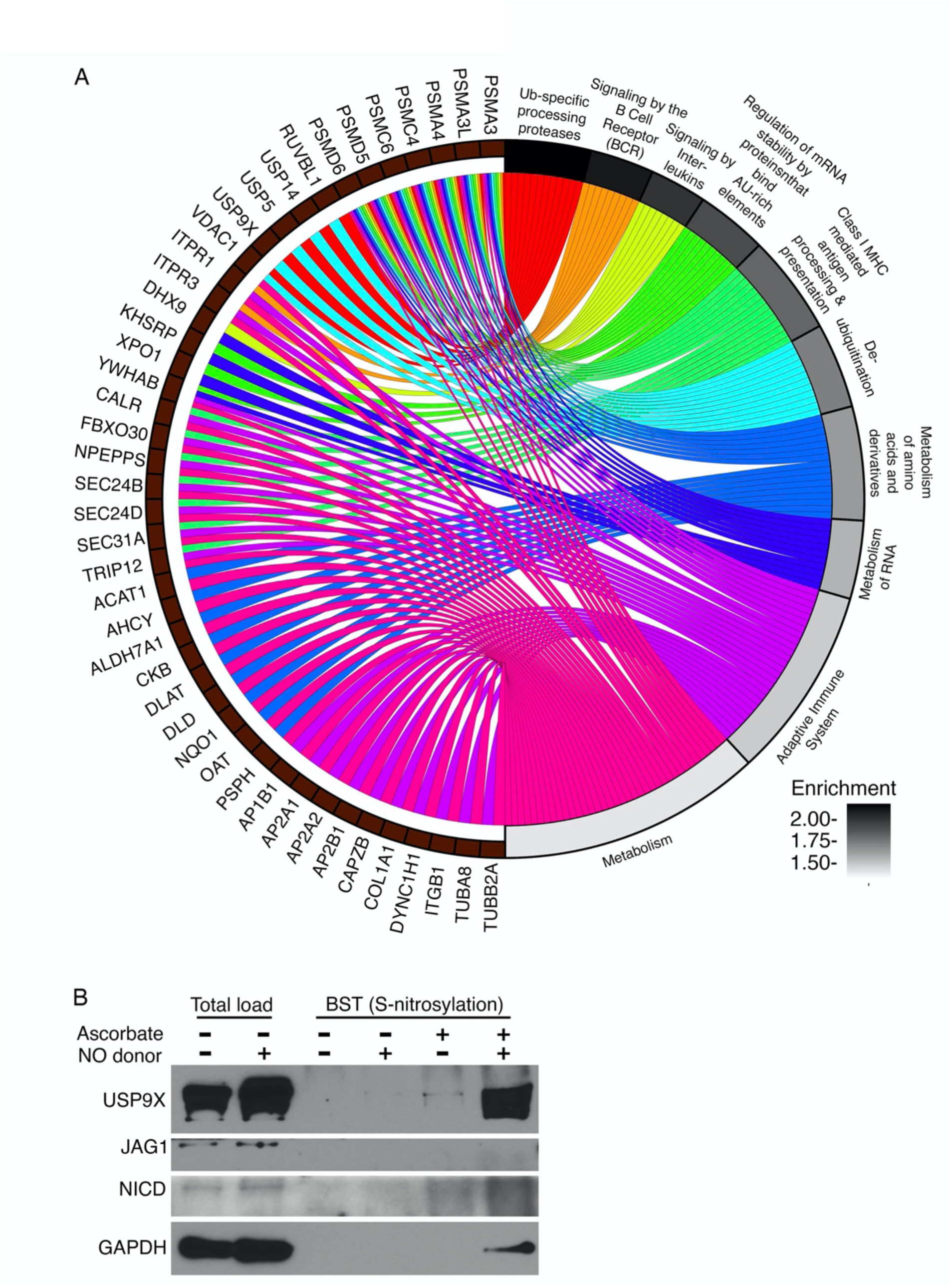
Mass spectrometric analysis of S-nitrosylated proteins in rat aortic valve interstitial cells (AVICs) exposed to NO donor demonstrate an enrichment of ubiquitin-specific processing proteases and identify USP9X. (A) Chord plot illustrates over-representation of NO-enriched S-nitrosylated proteins. Top 10 Gene Ontology pathways (GO Slim) and gene-sets are shown. (B) Immunoblot shows S-nitrosylation of USP9X in presence of NO donor, where negative ascorbate serves as a technical negative control. NICD and JAG1 immunoblots with ascorbate serve as biological negative control. GAPDH, a known S-nitrosylated protein, serves as a biological positive control. Total load represents the initial cell lysates utilized to purify S-nitrosylated proteins. BST, biotin switch technique.

Among the 12 S-nitrosylated proteins, we were specifically interested in USP9X (Ubiquitin specific peptidase 9, X-linked), given its previous link to NOTCH signaling^24^. *USP9X* is an evolutionarily conserved deubiquitinase and its Drosophila homolog, *faf*, regulates NOTCH signaling during eye development^25^. Further, in human breast cancer cell lines, USP9X deubiquitinates and stabilizes MIB1, an activator of ligand dependent canonical NOTCH1 signaling that is important for cardiac development^24,26^. Therefore, we postulated that nitrosylated USP9X deubiquitinates and stabilizes MIB1, which potentiates the ability of a ligand producing cell to activate NOTCH on a neighboring cell. First, we confirmed that USP9X undergoes post-translational modification by S-nitrosylation in pAVICs after NO treatment using modified BST followed by immunoblotting for USP9X (Figure 3B). In similar conditions, JAG1 and activated NICD, which are two important components of NOTCH1 signaling, did not show evidence of S-nitrosylation and served as negative controls. GAPDH, a well-known S-nitrosylated protein^27^, was used as a positive control (Figure 3B).

### Nitric oxide activates USP9X to deubiquitinate MIB1

To investigate if the deubiquitination activity of S-nitrosylated USP9X regulates NOTCH1, we examined the ubiquitination status of MIB1, a known activator of NOTCH1 signaling. For these studies, we generated stable HEK293 cells expressing both MIB1-FLAG and HA-Ubiquitin (Supplemental Figure 4A). This cell line expressing tagged MIB1 and Ubiquitin (Ub) was treated with NO donor, GSNO, or WP1130, a cell permeable small molecule reported as an inhibitor of deubiquitinases, including USP9X^28^ for 48 hours and proteasomal degradation was blocked by the addition of MG132 prior to the preparation of cell lysate. Immunoprecipitation of MIB1 using an anti-FLAG antibody was followed by measurement of HA-Ub-MIB1-FLAG using anti-HA antibody (Supplemental Figure 4A). For this experiment, S-nitrosylation of USP9X was confirmed after treatment with NO donor or GSNO while remaining at basal level in untreated control and after WP1130 treatment (Supplemental Figure 4B).

**Figure 4.**
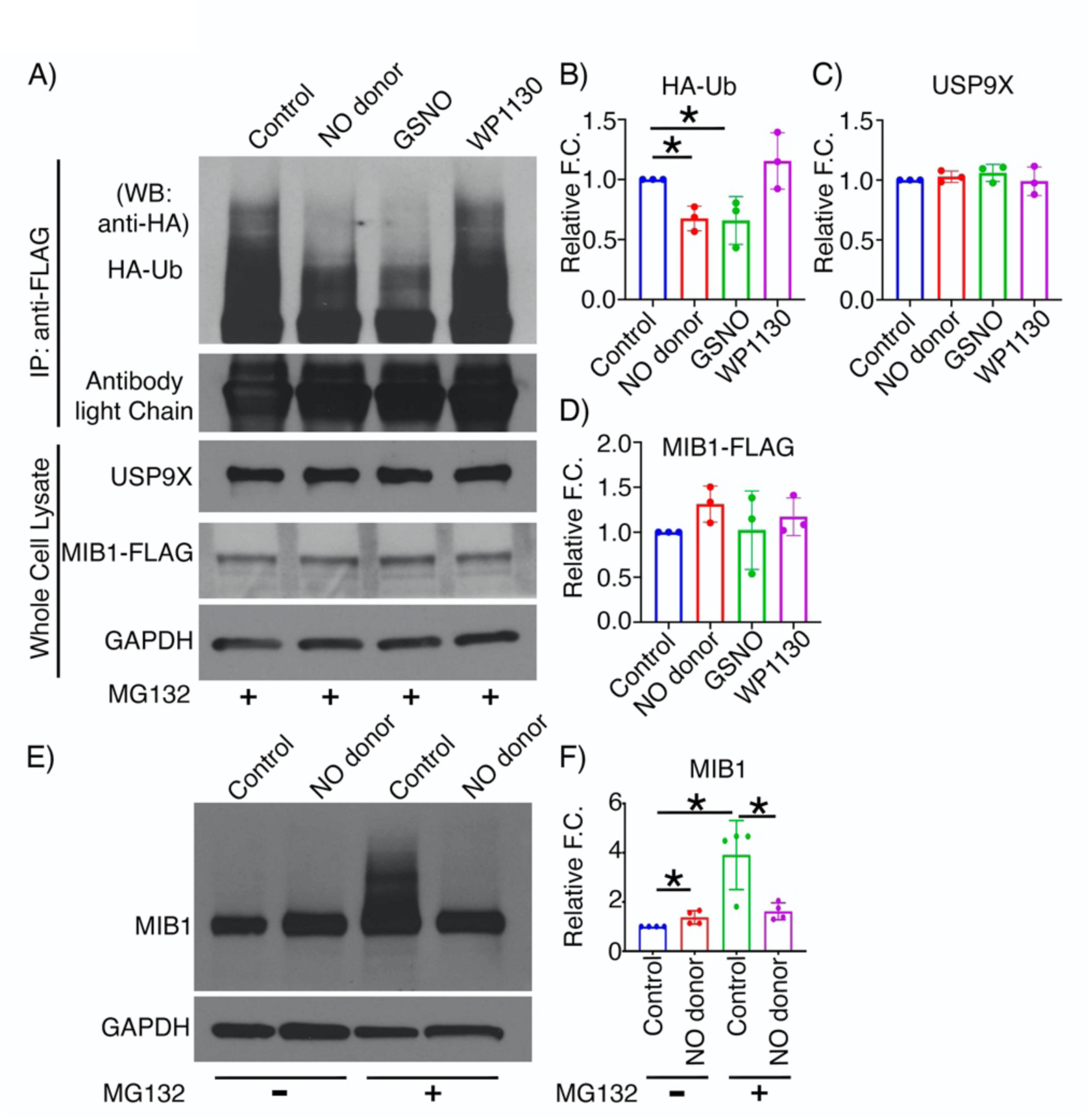
S-nitrosylation activates USP9X to deubiquitinate MIB1. HEK293 cells expressing MIB1-FLAG and HA-Ubiquitin (Ub) were cultured in the presence of detaNONOate (NO donor), GSNO (S-nitrosylating agent), or WP1130 (USP9X inhibitor). (A) After addition of MG132 and immunoprecipitation (IP) with anti-FLAG antibody, immunoblot with anti-HA antibody shows HA-Ub-MIB1-FLAG of various molecular weights in untreated condition (control) and with WP1130, indicating the presence of poly-Ub-MIB1. This is decreased with NO donor and GSNO. This is quantified in (B). (C, D) Quantification of USP9X and MIB1-FLAG expression after normalization against GAPDH. (E) Porcine AVICs were cultured in the presence and absence of NO donor for 5 days and immunoblot for MIB1 is shown with and without the addition of MG132. (F) Quantification of MIB1 is shown after normalization against GAPDH. * represent *P* value ≤ 0.05. For western blots (WB) 2-tailed unpaired t-test was performed. F.C., fold change.

Ubiquitin is an 8.6kD protein and varying degrees of ubiquitination are possible for a target protein (mono, multi-mono and poly-ubiquitination). Therefore, bands of increasing molecular weights are found by immunoblot after the proteasome block representing ubiquitination (Supplemental Figure 4A). With the addition of NO donor and GSNO, we found decreased ubiquitination of MIB1 when compared to treatment with deubiquitinase inhibitor, WP1130 or untreated control (Figure 4A-D). No significant difference in the expression of endogenous USP9X or MIB1-FLAG was observed, indicating upregulated deubiquitinase activity is due to S-nitrosylation (Figure 4A-D).

To examine the effect of S-nitrosylation of USP9X on the deubiquitination of MIB1, pAVICs were cultured for five days in presence and absence of NO donor. To measure ubiquitination, proteasomal degradation was inhibited by adding MG132, and ubiquitinated MIB1 was examined by immunoblot. We found decreased ubiquitination of MIB1 after NO donor treatment compared to the untreated condition, which confirms the NO-dependent activation USP9X in pAVICs (Figure 4E, F).

### Inhibition of USP9X reduces expression of MIB1 and NOTCH1 to promote calcification

To examine the effect of USP9X inhibition on NOTCH1 signaling and calcification of pAVICs, we utilized both pharmacological inhibition (with WP1130) and siRNA-mediated knockdown of USP9X (Figure 5). The addition of WP1130 to pAVICs decreased levels of MIB1 and NICD while treatment with NO donor was associated with increased MIB1 and NICD (Figure 5A, B; Supplemental Figure 5B, C). Accordingly, we also found increased RUNX2 expression with WP1130 treatment compared to untreated cells while RUNX2 expression decreased with NO donor (Figure 5C, D). Of note, culturing pAVICs with NO donor or WP1130 did not alter USP9X expression (Supplemental Figure 5A). In addition, our scRNAseq data indicated no significant alteration of *MIB1* expression at the transcriptional level after exposure to NO donor (Supplemental Figure 2B). Overall, these data indicate S-nitrosylated USP9X stabilizes MIB1, which is involved in NOTCH1 activation and inhibition of calcification.

**Figure 5.**
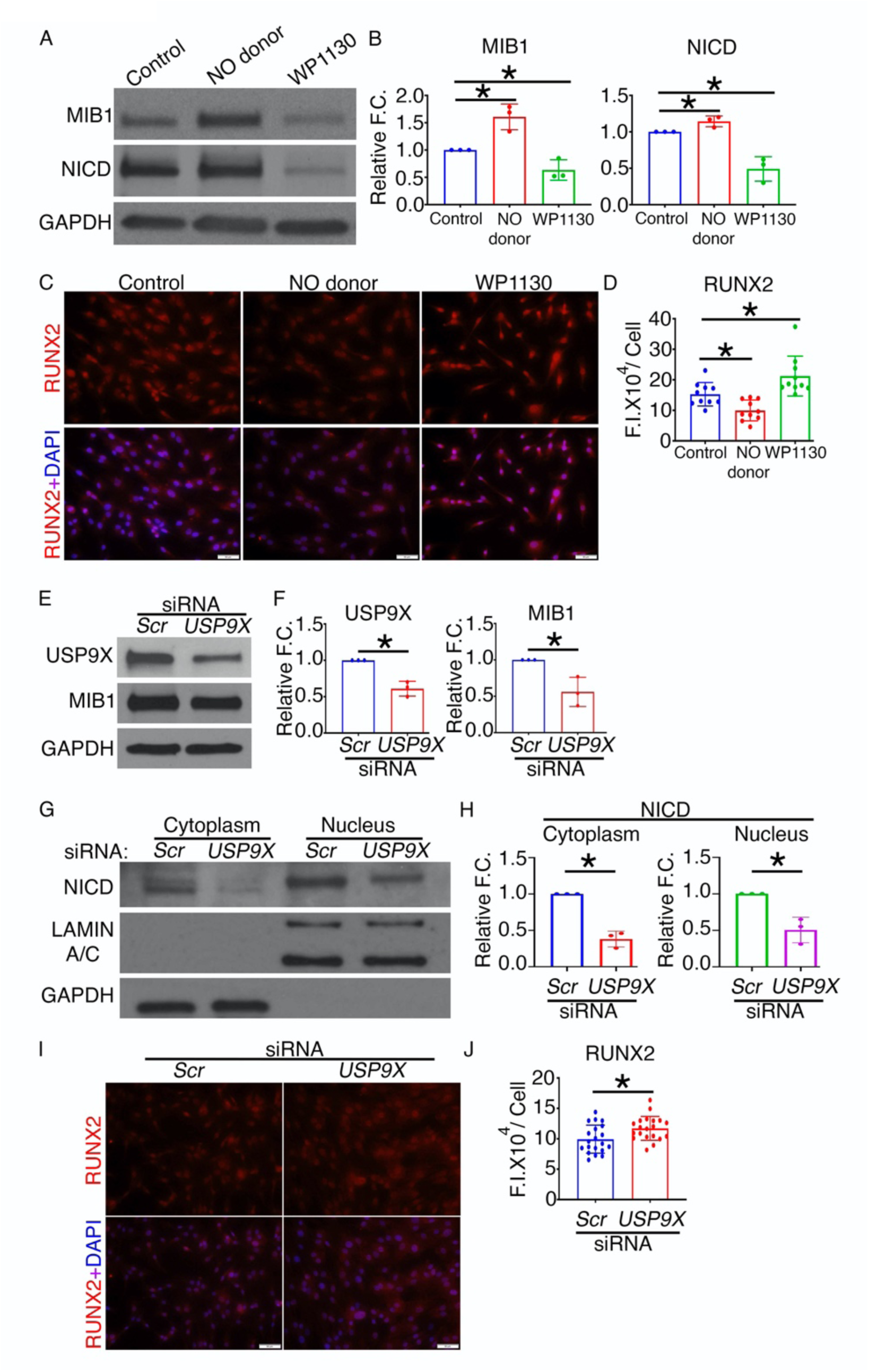
Inhibition of USP9X causes reduced expression of MIB1 and NOTCH1 intracellular domain (NICD) and increased RUNX2 expression in porcine aortic valve interstitial cells (AVICs). (A) Immunoblot shows decreased levels of MIB1 and NICD in porcine AVICs cultured with WP1130 for 5 days as compared to untreated cells (Control) while NO donor results in increased MIB1 and NICD. (B) Quantification of MIB1 and NICD expression is shown. (C) RUNX2 expression (red) is increased with WP1130 while RUNX2 is decreased with NO donor when compared to untreated cells (Control). Co-stained with nuclear DAPI (blue). (D) Quantification of RUNX2 expression is shown. (E) Addition of *USP9X*-siRNA to pAVICs reduced protein expression of USP9X and MIB1 as compared to scramble (*Scr*) control, which is quantified in (F). (G) With USP9x knockdown, nuclear NICD is reduced and this is quantified in (H) after normalization with LAMIN A/C (nuclear). Cytoplasmic NICD was normalized with GAPDH (G and H). (I) After treatment with *USP9X*-siRNA, RUNX2 expression (red) is increased when compared to scramble (*Scr*) control cells, co-stained with nuclear DAPI (blue). (J) Quantification of RUNX2 expression is shown. Scale bar: 50 μm. * represent p value ≤ 0.05 (2-tailed). For immunostaining and western blots, Mann-Whitney test and unpaired t-test were performed respectively. F.I.: Fluorescence Intensity; F.C.: Fold Change.

As WP1130 can target other deubiquitinases besides USP9X^28^, we utilized siRNA to knockdown *USP9X* for these studies. After optimization, we used 30nM concentration of pooled siRNA against *USP9X*, that resulted in a 60% reduction in *USP9X* mRNA and 40% reduction in USP9X protein (Figure 5E, F; Supplemental Figure 5D). With USP9X knockdown in pAVICs, we found reduced levels of MIB1 and NICD (Figure 5E-H; Supplemental Figure 5E-G). Additionally, we found evidence of osteogenic differentiation with upregulation of RUNX2 (Figure 5I-J; Supplemental Figure 5H, I). Altogether, these data indicate NO activates USP9X by S-nitrosylation, which deubiquitinates and stabilizes MIB1, and this is followed by the activation of NOTCH1 signaling to prevent calcification.

### Mice harboring endothelial cell-deletion of *Usp9x* display aortic valve stenosis and regurgitation

To determine if USP9X deficiency leads to AoV disease *in vivo*, we utilized mice in which *Usp9x* was genetically deleted. As germline deletion of *Usp9x* results in early embryonic lethality, mice harboring a floxed *Usp9x* allele (*Usp9x*^*fl/fl*^) were used that allowed condition deletion of the gene^29,30^. *Usp9x* resides on the X chromosome, therefore only male mice were used to test the effect of *Usp9x* deletion. *Usp9x*^*fl/fl*^ females were bred with males harboring *Tie2*^*Cre*^, which is expressed in the endothelial/endocardial cell lineage along with a subset of hematopoietic cells^31^ (Figure 6A). The progenies were genotyped at P10, and then only male mice were switched to a high fat western diet at 6 weeks of age. These mice subsequently underwent echocardiography at 6, 16 and 24 weeks of age followed by euthanasia to perform gross and cardiac histological examination (Figure 6B). We observed a partial lethality prior to P10 in *Usp9x*^*fl/Y*^; *Tie2*^*Cre*^ male mice, and is presumed to occur during embryogenesis or soon after birth (Figure 6C). Upon examination of the adult male survivors showed an increase in heart size within the *Cre*^*+*^ animals and confirmed reduction of USP9X protein expression in the AoVs of *Cre*^*+*^ mice as compared to *Cre*^-^ mice (Supplemental Figure 6A-B). Echocardiographic analysis of the *Cre*^*+*^ mice, revealed a significantly increased velocity across the AoV at 6, 16, and 24 weeks of age compared to *Cre*^-^ controls (*Usp9x*^*fl/Y*^; *Tie2*^*Cre-*^) indicative of functional AoV stenosis (Figure 6D). Additionally, an increase in left ventricular end systolic volume was noted at 16 and 24 weeks of age consistent with increased afterload associated with AoV stenosis (Supplemental Figure 6C). While not statistically significant, there was a trend towards decreased left ventricular ejection fraction and increased left ventricular end diastolic volume suggestive of ventricular dysfunction (Supplemental Figure 6C).

**Figure 6.**
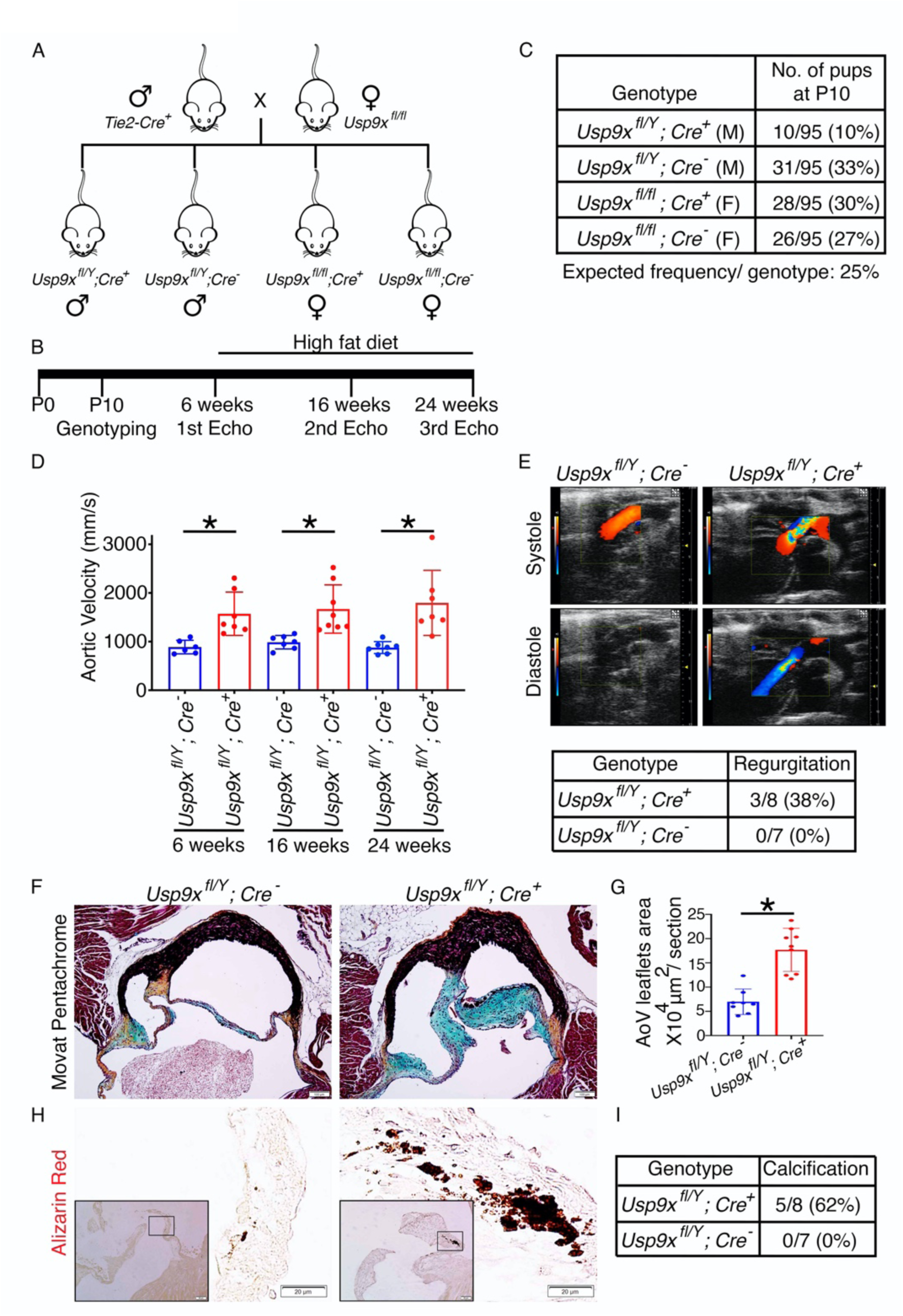
Aortic valve disease in mice with *Usp9x* deletion in endothelial cell lineage. (A) Breeding scheme to generate mice harboring endothelial and endothelial-derived cell deletion of *Usp9x* (*Usp9x*^*fl/fl*^; *Tie2*^*Cre*^). (B) Experimental scheme for *Usp9x*^*fl/Y*^; *Cre*^*+*^ males and *Usp9x*^*fl/Y*^; *Cre*^-^ controls. (C) Survival in *Usp9x*^*fl/Y*^; *Cre*^*+*^ male mice at postnatal day 10 (P10). (D) Aortic velocity measured by echocardiogram in *Cre*^*+*^ males at 6, 16 and 24 weeks of age compared to *Cre*^-^ controls. (E) Doppler image shows aortic valve regurgitation in *Cre*^*+*^ male at 24 weeks of age as compared to *Cre*^-^ control. Table shows incidence of aortic valve regurgitation in *Cre*^*+*^ and *Cre*^-^ males. (F) Histological section of aortic valve stained with Russell-Movat pentachrome shows increased proteoglycan deposition in *Cre*^*+*^ males compared to *Cre*^-^ controls. (G) Graph shows quantification of aortic valve area in *Cre*^*+*^ and *Cre*^-^ males. (H) Alizarin-Red staining demonstrates calcification in *Cre*^*+*^ males. (I) Table shows incidence of calcification in *Cre*^*+*^ and *Cre*^-^ males. Scale bar: Pentachrome staining 100 μm; Alizarin red staining 20 μm. * represent p value ≤ 0.05 (2-tailed Mann-Whitney test).

Three out of 8 *Cre*^*+*^ mice (38%) demonstrated AoV regurgitation, indicative of incomplete coaptation of the AoV leaflets compared to *Cre*^-^ (0/7) controls (Figure 6E, Online Video I, II). Histological analysis using Russell-Movat Pentachrome staining of AoVs collected from 6-month-old mice revealed proteoglycan accumulation throughout the thickened leaflets of *Cre*^*+*^ mice as compared to *Cre*^-^ control (Figure 6F, G). Evidence of calcification was detected by positive Alizarin Red staining in 5 out of 8 *Cre*^*+*^ mice (62%) as compared to the lack of staining in the valves of *Cre*^-^ control mice (0/7) at 6 months of age (Figure 6H, I). Expression of RUNX2 was increased, while MIB1 and NICD were decreased in *Cre*^*+*^ as compared to *Cre*^-^ control mice (Supplemental Figure 6D, E).

### Calcified human aortic valves display reduced S-nitrosylation of USP9X

To determine if the S-nitrosylation of USP9X was altered in CAVD, we obtained explanted AoV tissue from patients at the time of surgical valve replacement. Aortic stenosis was the indication for surgery, and all of the patients had tricuspid aortic valves. The disease was categorized as moderate (n=2) or moderate-severe (n=2) valve calcification (Supplemental Figure 7A). In all calcified valve leaflets, RUNX2 expression was increased, as expected (Figure 7A). We also found a reduction in the amount of S-nitrosylated USP9X, which was inversely correlated with increased disease severity (Figure 7A). Furthermore, there was reduced expression of MIB1, NICD and the Notch ligand, JAG1, when compared to a control non-calcified aortic valve (Figure 7A; Supplemental Figure 7B). Of note, we also found the overall expression of USP9X protein reduced in calcified valves (Figure 7A).

**Figure 7.**
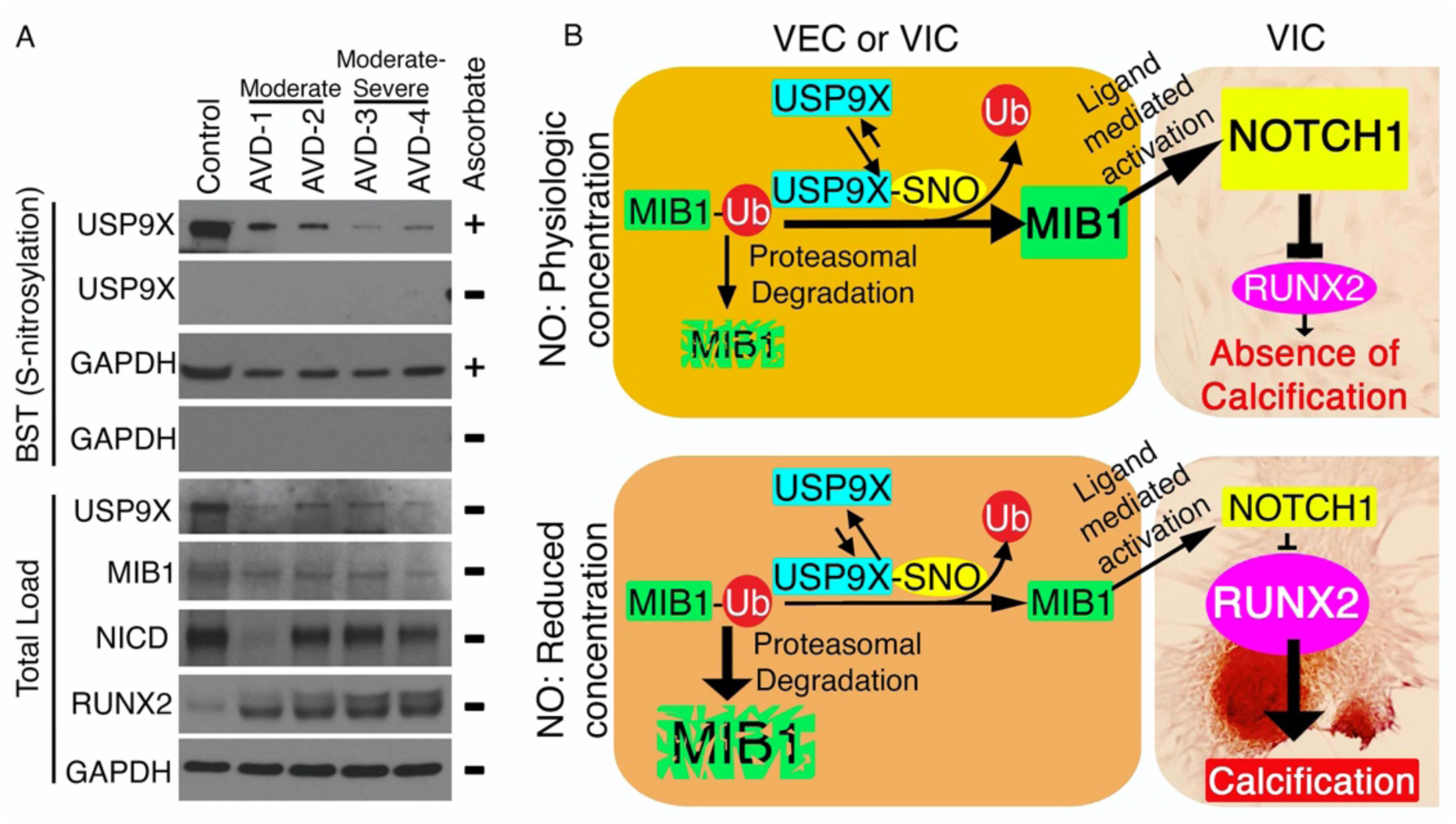
Calcified human aortic valves display reduced S-nitrosylation of USP9X along with decreased expression of MIB1 and NOTCH1 intracellular domain (NICD). (A) Immunoblot for USP9X shows significantly reduced S-nitrosylation in aortic valves from adults with moderate (AVD-1 and 2) and moderate-severe (AVD-3 and 4) calcific aortic valve disease when compared to uncalcified aortic valve (Control). S-nitrosylation of GAPDH is also shown. Absence of a band in ascorbate negative experiment served as the negative control. Total load shows protein expression in the cell lysates prior to purification of S-nitrosylated proteins where USP9X, MIB1 and NICD expression is reduced in calcified valves while RUNX2 is increased. GAPDH serves as a loading control. (B) Schematic showing proposed model where S-nitrosylation of USP9X leads to MIB1 stabilization via deubiquitination, which increases ligand-mediated NOTCH1 activation in neighboring cells and the inhibition of calcification in physiological concentrations of NO. This pathway is abrogated under low concentrations of NO.

Overall, these data demonstrate NO-mediated inhibition of AoV calcification via activation of a USP9X-MIB1-NOTCH1 signaling axis. We propose that under physiological NO concentrations, S-nitrosylation of USP9X results in stabilization of MIB1 by its deubiquitination, which prevents proteasomal degradation (Figure 7B). The stable MIB1 subsequently triggers ligand-dependent NOTCH1 activation in neighboring cells, which inhibits RUNX2 expression and the calcification gene program. At low NO concentrations, USP9X is not activated by S-nitrosylation allowing for degradation of MIB1 through the ubiquitin-proteasome pathway. The loss of MIB1 prevents ligand-dependent NOTCH1 activation thereby promoting calcification (Figure 7B).

## Discussion

In this study, we demonstrated a novel molecular mechanism involving the deubiquitinase, USP9X, by which NO regulates NOTCH1 signaling in the setting of CAVD. Here, we show for the first time that S-nitrosylation, a NO-dependent posttranslational modification, is important for the prevention of CAVD through inhibition of myofibroblast activation of AVICs. Our proteomic analysis revealed the role of S-nitrosylation in regulation of the ubiquitin proteasome pathway. We also discovered that S-nitrosylation of USP9X is critical for its stabilization of MIB1, which potentiates the ligand mediated activation of NOTCH1 signaling. Furthermore, deletion of *Usp9x* in the endothelial cell lineage in mice resulted in aortic valve stenosis and calcification. Finally, translational studies using calcified human AoVs showed alterations in levels of S-nitrosylated USP9X, MIB1 and NOTCH1 intracellular domain that were consistent with our proposed mechanism (Figure 7B).

Our group along with others have reported the protective role of a healthy valve endothelium in CAVD, and in our previous publication showed that this protection is partly achieved by endothelial-derived NO regulation of NOTCH signaling^10,18,32^. Using an ex vivo approach, a protective effect from endothelial NO that was mediated by a sGC/cGMP-dependent pathway was also reported by Richards et al.^18^. A sGC/cGMP-dependent mechanism has also been suggested by the investigation of mechanisms of vascular calcification where NO was shown to regulate TGF-β signaling^33^. While we did not find that activation of sGC/cGMP was sufficient to protect pAVICs from calcification in our *in vitro* system, it remains to be determined if elevated NO production by increased laminar blood flow or shear stress with subsequent activation of sGC/cGMP-pathway is protective against CAVD.

Apart from the NOTCH pathway, USP9X can also regulate the Wnt, TGF-β and Hippo signaling pathways, which are also important for AoV development and disease^29,34^. The impact of S-nitrosylation of USP9X on these pathways is not known. In addition to USP9X, we also have identified S-nitrosylation of other ubiquitin pathway members in our proteomic analysis of AVICs treated with NO donor and they include USP5 and USP14 (Figure 3A). Similar to USP9X, USP5 and USP14 are also reported to regulate NOTCH and Wnt signaling pathways in a variety of cancers and also in Drosophila eye development^35-37^. USP5 also regulates EMT by stabilizing SLUG, while USP14 controls ciliogenesis by regulating Hedgehog signaling^38,39^. Both of these pathways have well-established roles in valve morphogenesis but have not been investigated in valve calcification^40,41^. Intriguingly, the ubiquitin proteasome pathway was the topmost, and the deubiquitination pathway was in top ten pathways in our proteomic analysis (Figure 3A). The ubiquitin proteasome pathway, including deubiquitination, has already drawn attention as a potential drug target for other diseases, such as cancer, neurodegenerative diseases, and immunological disorders as it is involved in numerous key regulatory processes. The small number (<100) of deubiquitinases are likely subject to multiple layers of regulation that modulate both their activity and their specificity^42^. However, the role of the ubiquitin-dependent pathway in valve development and disease, including calcification, has not been studied. To the best of our knowledge, this is the first report demonstrating the association between the ubiquitin pathway and valve calcification.

Our data suggests the S-nitrosylated USP9X deubiquitinates to stabilize MIB1, which then ubiquitinates the NOTCH-ligands; a crucial step for ligand internalization and NOTCH1 activation in neighboring cells^43,44^. The importance of MIB1 and JAG1 for AoV development and calcification has been shown. Deletion of *Mib1* in cardiac progenitor cells utilizing *Nkx2*.*5-Cre*, led to neonatal lethality due to valve dysfunction and deletion of *Jag1* in endothelial cells using a *Cdh5-Cre* resulted in a spectrum of cardiac phenotypes from congenital heart defects reminiscent of tetralogy of Fallot to adult valve disease characterized by thickened semilunar valves and calcification^26 45^. We utilized *Tie2-Cre* mice to delete *Usp9x* in endothelial and endothelial-derived cells and observed an embryonic/perinatal lethality of ∼58% (Figure 6C). The etiology of this lethality is under investigation.

As both *Cdh5* and *Tie2* express prior to EMT, these *Cre* lines can delete target genes in endothelial and endothelial-derived cells, such as VICs^31^. Therefore, we are unable to determine whether USP9X function is important either in VECs, VICs or both. In an attempt to address this question *in vivo*, we deleted *Usp9x* utilizing *Twist2-Cre*, which can delete target genes in only VICs^46^. However, we were unable to obtain any live *Usp9x*^*fl/Y*^*;Twist2*^*Cre+*^ mice after P10 precluding its study in CAVD (data not shown). We also performed *in vitro* experiments using pAVECs and pAVICs. Here, *USP9X* knockdown was performed using siRNA against *USP9X* in pAVECs, pAVICs, or both. When pAVECs or pAVICs were cultured alone, we found increased expression of RUNX2 in pAVICs at baseline and *USP9X* knockdown in either cell type resulted in increased expression, although it was more dramatic in pAVICs (Supplemental Figure 8A-D, I). When pAVICs and pAVECs were cocultured, we observed decreased RUNX2 expression, as expected (Supplemental Figure 8C, E, I). In this coculture system, *USP9X* knockdown in either pAVICs or pAVECs also resulted in increased RUNX2 expression but here it was more dramatics in pAVECs (Supplemental Figure 8E-I). These *in vitro* experiments support roles for USP9X in both valve endothelial and interstitial cells in CAVD.

Our scRNA-seq analysis of pAVICs, which spontaneously calcify *in vitro*, and reports from other labs demonstrate that VICs in culture conditions are already activated towards myofibroblast differentiation with the expression of the stress fiber gene, *ACTA2* ^21,47^. This myofibroblast activation can be inhibited by culturing VICs into 3D hydrogel or by using a fibroblast media formulation with collagen coated surface that maintain a quiescent phenotype of VICs^21,47^. Here, we observed that S-nitrosylation can reverse myofibroblast-VICs towards an inactivated/quiescent VICs by reducing expression of *ACTA2, VIM, TAGLN* and *CNN1* (Figure 2B). However, this phenotypic reversal is temporary, and always requires elevated NO concentration in culture, which indicates that S-nitrosylation is required for maintenance of the quiescent state of VICs (Figure 2C). The *in vivo* significance of this observation for CAVD pathogenesis requires further study.

In a healthy valve, NO is primarily produced by NOS3, which is expressed by endothelial cells^20^. Expression of the *NOS3* gene is regulated by the transcription factors, KLF2 and KLF4. KLF2 and KLF4 are known to be regulated in endothelial cells by blood flow via the mechanosensor, PLEXIND1^48^. Flow-dependent endothelial cell-derived NO is a freely diffusible, but short-lived molecule. Its diffusion coefficient in tissue is nearly fourfold smaller than in solution^49^. This low diffusion rate may restrict the ability of NO to travel from VECs of the ventricular side to the VICs populating the aortic side of the leaflet. We observed increased S-nitrosylation on the flow side of the aortic and mitral valves in mice (Supplemental Figure 8J). Therefore, an increased amount of activated USP9X and NOTCH1 signaling are expected in the ventricular side of the aortic valve, which is consistent with the observation that calcific nodules primarily occur on the fibrosa/aortic side of the human aortic valve^50^. Notably, mouse heart valves are much thinner compared to human heart valves and therefore the bioavailability of diffusible NO is more in mice as compared to humans. This may explain the relative difficulty in developing models of CAVD in mice, and potentially explains why additional genetic manipulation that results in thickened leaflets is required to see the development of CAVD in *Notch1* heterozygous mice^10,51^.

In conclusion, we have discovered a novel molecular mechanism by which endothelial NO regulates the NOTCH1 signaling pathway in CAVD that involves the ubiquitin-proteasome system. These findings open the door to investigate regulators of the ubiquitin-proteasome pathway as potential therapeutic targets in CAVD.

## Methods

### AVECs and AVICs culture and treatments

Porcine aortic valve interstitial cells (pAVICs) and endothelial cells (pAVECs) were collected from juvenile pig valve cusps and isolated as previously described^52^. Briefly, valve leaflets were subjected to 5 mins collagenase (Worthington Biochemical# LS004176) digestion at 37°C and the endothelial layer was gently sheared by sterile swab onto the leaflet. Dislodged pAVECs were collected from the cell suspension and cultured in endothelial media as described earlier^52^. At near confluency, the pAVECs were passaged with TrypLE Express (Fisher Scientific# 12-604-021) and further cultured in EGM-2 media (Lonza# CC-3162). pAVECs used in this study were between passages 2 and 4. After removal of the endothelial layer, valve leaflets were subjected to a 15-hour collagenase digestion at 37°C and cultured the pAVICs in an interstitial porcine medium as previously described^52^. At near confluency, pAVICs were passaged with trypsin-EDTA and maintained in Media-199 (ThermoFisher# 11150059). pAVICs used in this study were between passages 3 and 7. Rat aortic valve interstitial cells (rat-AVICs) were similarly isolated from Sprague-Dawley adult rats by a collagenase digestion and cultured in Media-199. To identify S-nitrosylated proteins, we used the modified biotin switch technique (BST) followed by mass spectrometry (MS). Rat-AVICs were cultured in osteogenic media, by supplementing Media-199 with 50μg/ml ascorbate-2-phosphate (Sigma-Aldrich# 49752), 10nM dexamethasone (Sigma-Aldrich# D4902) and 10μM β-glycerol phosphate (Sigma-Aldrich# G9422). For each cell culture experiments performed in presence of exogenous reagents, media were supplemented with either detaNONOate (NO donor: 150μM) (FisherScientific# AC328651000), or S-nitrosoglutathione (GSNO) (S-nitrosylating agent: 200μM) (MilliporeSigma# N4148), or 8-Br-PET-cGMP (cGMP analog: 0.025μM) (Axxora# BLG-P003-10), or WP1130 (Usp9x inhibitor: 1μM for HEK293 and 0.05μM for pAVICs) (Cayman# 15227) and were refreshed daily. To stop proteasomal degradation, 10μM MG132 (MilliporeSigma# 474787) was added to the culture media 10 hours before preparation of the cell lysate.

### Alizarin Red staining

To stain calcific nodules, pAVICs were fixed in 2.5% PFA for 15 minutes at 4°C, incubated with Alizarin Red Solution (Millipore# 2003999) for 5-7 minutes, followed by washing with distilled water. The number of nodules per well were counted and images were captured using Olympus BX51 microscope attached with Olympus DP71 camera.

### Immunofluorescence staining of S-nitrosylated proteins

To visualize the S-nitrosylated proteins in pAVICs (n≥3) and in three months old murine valve tissue section, biotin switch technique was performed using S-Nitrosylated Protein Detection Kit (Cayman Chemical# 10006518). Immunofluorescence staining was performed following the manufacturer’s protocol and images were captured using Olympus IX51 microscope attached with Olympus DP72 camera. Nuclei were stained with DAPI.

### DAR4M staining

To estimate endogenous levels of NO, unfixed pAVICs were incubated with 10μM DAR-4M AM (Sigma-Aldrich# 251765) for 15 minutes at room temperature, followed by washing with PBS. Nuclei were stained with DAPI. Images were captured using the Olympus IX51 microscope attached with Olympus DP72 camera.

### Single cell RNA sequencing and analyses

pAVICs were dissociated with TrypLE Express, followed by filteration through a 40μm cell strainer (BD Falcon# 352340), and resuspension in 0.04% BSA. Cell viabilities (>85%) were measured using the Automated Cell Counter (ThermoFisher# AMQAF1000). Single-cell droplet libraries from this suspension were generated in the 10XGenomics Chromium controller (∼2000 target cell recovery/group) according to the manufacturer’s instructions. For each sample, the quality and integrity of cDNA and cDNA libraries were quantitated using the High Sensitivity D5000 and D1000 ScreenTape↓ (Agilent# 5067-5592 and 5067-5584) on the Agilent-2200 TapeStation. All libraries were sequenced (150bp paired-end) in Illumina HiSeq4000 platform at The Steve and Cindy Rasmussen Institute for Genomic Medicine in Nationwide Children’s Hospital.

We used the 10X Genomics’ Cell Ranger pipeline to demultiplex and convert Illumina bcl output into fastq files. We mapped the fastq reads to the pig genome Sscrofa11 with Y sequences from WTSI_X_Y_pig V2 (GCF_000003025.6_Sscrofa11.1_genomic.fna) and gene annotation (GCF_000003025.6_Sscrofa11.1_genomic_genes.filtered.gtf) downloaded from NCBI (https://www.ncbi.nlm.nih.gov/assembly/GCF_000003025.6/). scRNA-seq data with different treatments (control, GSNO, NO donor and NO donor withdrawal after three days) were combined using the cell ranger aggregate function yielding a total of 7037 cells (control: 1683, GSNO: 1966, NO donor: 1485 and NO donor withdrawal: 1903) with an average of 3397 genes per cell. This data was processed using Seurat (version 3.0)^53^ to perform normalization and identification of genes with the highest variability. Top 100 genes that showed significant differential expression (adjusted P Value ≤ 0.05) between control and NO treated samples were used to plot the heatmap. Z-score transformed scaled expression of genes between the four samples were plotted using DoHeatmap function of Seurat (3.0) package in R.

Genes upregulated (green) and downregulated (pink) are shown in a volcano plot based on negative log_10_P value (2 cut-off) between control and NO treated samples (Supplementary Figure 2). Volcano plot was created using ggplot2 (3.2.1) package of R.

In parallel, we imported P30 mouse aortic valve data^20^ from GEO omnibus database (GSE117011). To compare the mouse data to our porcine data, we replaced the mouse gene names with corresponding porcine homolog’s gene names downloaded from BioMart (Ensemble; https://useast.ensembl.org/info/data/biomart/index.html). Mouse gene names without a clear porcine homolog were retained as mouse gene names. This data was also processed using Seurat similar to the porcine single-cell data. The mouse and the porcine data were then integrated, scaled, and dimensionality reduction was performed using Seurat. Violin plots were created to show marker-gene expression in different clusters and drawn using a modified VlnPlot from Seurat (3.0) package in R.

Differential gene expression analysis was performed by splitting the pAVICs of one treatment into three sample groups of equal number of cells. These *in silico* replicates were then analyzed using DESeq2^54^ to identify differentially expressed genes between conditions. We compared the gene expression changes in the pAVICs to the expression changes observed in human AVECs subjected to shear stress^22^ by comparing the relative gene expression changes in each sample. For the pAVICs, we used relative changes with respect to control cells and for the human AVECs, we used control cells treated with shear stress as our baseline. Shear responsive genes with a significant impact (*P* value ≤ 0.05) in their expression in endothelial cells after *NOTCH1*-knockdown using siRNA, were used to identify homologs in the porcine genome. Genes that had clear homologs in porcine genome (downloaded from BioMart), and those that were expressed in pAVICs (at least 1 read count in any of the samples), were evaluated for changes in expression following NO treatment.

### BST and mass spectrometry

Biotin switch technique (BST) was performed as previously described with minor modifications^55^. Until the purification of the S-nitrosylated proteins, all of the procedures were performed in the dark. Briefly, cultured cells or frozen aortic valve tissues that were crushed under liquid nitrogen were harvested in HEN (100mM HEPES, 1mM EDTA, 0.1mM Neocuproine) lysis buffer containing 50mM NaCl, 0.2% methyl methanethiosulfonate (MMTS), 1mM PMSF, 1% NP-40, 1tab/10ml protease inhibitor cocktail (MilliporeSigma# 11836170001). Protein concentrations in the cell lysates were estimated using the Quick Start™ Bradford Dye Reagent (Biorad# 5000205) followed by blocking of free thiol groups in HEN buffer containing 0.2% MMTS and 2.5% SDS at 50°C for 30 minutes. After blocking, the proteins were purified by acetone precipitation and further resuspended in HENS (HEN+1%SDS) buffer. These proteins were designated as “total load”. Further purification of S-nitrosylated proteins were done using Thiopropyl Sepharose 6B bead (GE Healthcare# 170420001) in the presence of 50mM Na-ascorbate as a mild reducing agent, which will reduce -SNO to -SH, keeping the disulfide bond (-S-S-) intact. For the negative control, Na-ascorbate was replaced by HENS buffer. Purified proteins were eluted in SDS-PAGE sample loading buffer and separated in 4-20% Mini-PROTEAN^®^ TGX™ Precast Gels (Bio-Rad# 4561094, 4561096) either for western blot (WB) or stained with Imperial™ Protein Stain (Thermo Scientific# 24615). No band in the negative control lane demonstrated specificity of the purified S-nitrosylated proteins (Supplemental Figure: 3B). Stained lanes were excised from the gel and sent for mass spectrometry (LC/MS-MS) at the Center for Proteomics and Bioinformatics, Case Western Reserve University. MassMatrix bioinformatics software^56^ was used to search the acquired MS data against rat database from Uniprot. Identified S-nitrosylated proteins with 0% decoy were selected for analysis. We performed a total of 6 mass spectrometry runs, triplicate for both with and without NO donor. S-nitrosylated proteins were designated if present in at least two out of six samples. Any protein identified in more than 2 samples in the presence of NO donor, compared to samples in the absence of NO donor were designated as NO-enriched. NO-enriched proteins were analyzed utilizing WebGestalt 2019 software (WEB-based Gene SeT AnaLysis Toolkit)^23^ to identify over-represented pathways and associated proteins. Top 10 GO (gene ontology) Slim pathways and associated proteins were used to create the chord plot using a modified GOChord script in R.

### Generation of stable cell lines

HEK293 cells were transfected with pMIB1-FLAG (Addgene# 37116) and pHA-Ub (Addgene# 18712) plasmids using Lipofectamine 3000 (ThermoFisher# L3000015) following the standard protocol. Stable cells were selected by culturing transfected cells in DMEM (ATCC# 30-2002) media in the presence of 500μg/ml Geneticin (ThermoFisher# 10131035) and 10μg/ml Puromycin (ThermoFisher# A1113803). For MIB1 ubiquitination assay, transfected HEK293 cells were cultured in DMEM media without antibiotics for 3 days.

### *USP9X* knockdown by siRNA

A pool of three target-specific (19- to 25-nucleotide) siRNAs was used to silence *USP9X* expression (Supplemental Table 2). The siRNAs targeting porcine *USP9X* (ENSSSCG00000012251) transcript were designed using BLOCK-iT™ RNAi Designer (ThermoFisher Scientific) and synthesized from Eurofins Genomics. A control siRNA was used as a scramble (*Scr*) siRNA, targeting no known gene. pAVICs maintained at 60%– 70% confluency was transfected with 30nM *USP9X* or *Scr* siRNA using Lipofectamine 3000 Transfection Reagent (L3000015, ThermoFisher Scientific). Cell lysates were collected and examined for USP9X protein expression 72 hours post-transfection.

For co-culture experiments, pAVECs and pAVICs were transfected separately. After 12 hours of transfection, cells were detached using TrypLE Express and co-cultured in EGM-2 media (Lonza# CC-3162) in different combinations for additional 60 hours as described in Supplemental Figure 8A-H.

### RNA purification and quantitative real-time PCR

RNA was extracted from pAVICs using TRIzol Reagent (ThermoFisher Scientific# 15596018) following the manufacturer’s instructions. 0.5-1.0μg of RNA was used for reverse transcription using SuperScript VILO cDNA Synthesis Kit (ThermoFisher Scientific# 11754-050). SYBR Green–based quantitative real-time PCR was performed using the Applied Biosystems 7500 real-time PCR. Mean relative gene expression was calculated after normalization of C_t_ values to *GAPDH* using the ^ΔΔ^Ct method (Supplemental Table 1).

### Immunoprecipitation and western blot analysis

Cell lysates were prepared from cultured cells in an assay specific buffer. For immunoprecipitation (IP), the cell lysate was prepared in Pierce™ IP Lysis Buffer (ThermoScientific# 87787) and for western blot (WB) in RIPA Lysis and Extraction Buffer (ThermoFisher Scientific# 89900) supplemented with Halt Protease Inhibitor Cocktail (ThermoFisher Scientific# 87785). Cell lysates were centrifuged at 15,871g for 15 minutes at 4°C and the supernatants were collected. Protein concentrations were estimated using the Pierce BCA Protein Assay Kit (ThermoFisher Scientific# 23227). 400-500μg of protein from the stably transfected HEK293 cells were used to immunoprecipitate FLAG-tagged MIB1 using EZview™ Red ANTI-FLAG® M2 affinity gel (MilliporeSigma# F2426) following the manufacturers protocol at 4°C overnight.

For nuclear-cytoplasmic fractionation, NE-PER™ Nuclear and Cytoplasmic Extraction Reagents (Thermo Scientific# 78833) were used following the manufacturer’s protocol. To test the purity of nuclear and cytoplasmic extracts, a WB against LAMIN A/C (Santa Cruz Biotechnology# sc-7292) and GAPDH (Novus Biologicals# NB300-221) were performed respectively.

For the WBs 10-25μg of the protein samples were boiled for 5 minutes with 6X Laemmli SDS-Sample Buffer (Boston BioProducts# BP-111R) containing β-mercaptoethanol and separated in 4-20% Mini-PROTEAN^®^ TGX™ Precast Gels (Bio-Rad# 4561094, 4561096). After separation, the proteins were transferred into a polyvinylidene difluoride membrane (Bio-Rad# 1620177), and blocked with 5% nonfat milk in TBS containing 0.1% Tween®-20. Membranes were probed with primary antibodies against USP9X (Cell Signaling Technology# 5751), MIB1 (Santa Cruz Biotechnology# sc-393811), NICD (Abcam# ab8925 and Cell Signaling Technology# 4147), JAG1 (Santa Cruz Biotechnologies# sc8303 and Santa Cruz Biotechnologies# sc390177), RUNX2 (Cell Signaling Technology# 8486), HA-tag (Sigma-Aldrich# H-9658), FLAG (Sigma-Aldrich#), phospho-VASP (Cell Signaling Technology# 3114). After probing with primary antibodies, membranes were further probed with HRP-conjugated anti-rabbit and anti-mouse secondary antibodies (Vector Laboratories# PI-1000, PI-2000). Immunoblots were developed using either Pierce ECL Western Blotting Substrate (ThermoFisher Scientific# 32106) or SuperSignal West Dura Extended Duration Substrate (ThermoFisher Scientific# 34075). Densitometric analysis was performed to normalize the protein levels with respect to GAPDH or otherwise mentioned as loading control using ImageJ software. For re-probing with different primary antibodies, membranes were stripped with Restore™ Western Blot Stripping Buffer (Thermo Scientific# 21059) following the manufacturer’s protocol.

### Immunofluorescence and immunohistochemistry in vitro and in vivo

For immunofluorescence (IF) staining, cultured pAVICs were fixed in 2.5% PFA. Cells were permeabilized with PBST (PBS containing 0.1% TritonX100), and nonspecific immunoreactions were blocked using 1% BSA in PBST for 1 hour at room temperature. pAVICs were incubated overnight with primary antibodies against RUNX2 (Abcam# ab76956), activated NICD (Abcam# ab8925), followed by washing with PBST and incubating with Alexa Fluor-594 conjugated anti-rabbit secondary antibody (ThermoFisher Scientific# A21207) for 1 hour in the dark. Nuclei were stained with 1.5ug/ml DAPI (Sigma-Aldrich# D9542). Images were captured using Olympus IX51 microscope attached with Olympus DP72 camera.

A similar protocol was performed to stain the tissue sections. The section underwent deparaffinization, using xylene and grades of ethanol, followed by antigen retrieval using citrate-based Antigen Unmasking solution (Vector laboratories# H-3300) following the manufacturer’s protocol. After permeabilization and blocking with 1% BSA for 1 hour, tissue sections were incubated with primary antibodies against MIB1 (Abcam# ab124929), USP9X (Cell Signaling Technology# 14898) and RUNX2 (Abcam# ab76956) overnight at 4°C. Following PBST wash, sections were incubated with anti-rabbit and anti-mouse secondary antibodies that were conjugated to Alexa Fluor 594/568/488 (ThermoFisher Scientific# A21207, A10037, A11001, A11008) for 1 hour at room temperature. Nuclei were stained with VECTASHIELD® HardSet™ Antifade Mounting Medium with DAPI (Vector Laboratories# H-1500). Images were visualized using Olympus BX51 microscope attached with Olympus DP71 camera.

For immunohistochemical (IHC) staining, tissue sections were deparaffinized in xylene and rehydrated in grades of ethanol and PBS, followed by an antigen retrieval. Sections were incubated at room temperature with 3% H_2_O_2_ for 10 minutes to quench endogenous peroxidase activity and blocked by 5% normal goat serum (Vector Laboratories# S-1000) in TBST (TBS containing 0.1% Tween® 20) for 1 hour. After blocking, sections were incubated with a primary antibody against NICD (Abcam# ab8925) overnight at 4°C. Following primary antibody incubation sections were further incubated with SignalStain Boost anti-Rabbit IHC Detection Reagent (Cell Signaling Technology# 8114) at room temperature for 30 minutes. Sections were visualized utilizing SignalStain DAB Substrate Kit (Cell Signaling Technology# 8059) and imaged using Zeiss AxioImagerA2 attached with AxioCam MRc camera.

### Experimental Mouse Model

Animal experiments were approved by the Institutional Animal Care and Use Committee at the Research Institute at Nationwide Children’s Hospital. *Usp9x*^*fl/fl*^ mice were obtained from Charles River Laboratories, Italy, originally made by *Ozgene Pty Ltd*, Australia with permission of Dr. Stephen Wood, Griffith University, Australia^30^. *Usp9x*^*fl/fl*^ females were bred with*Tie2*^*Cre*^ males (Jax# 008863), to obtain experimental *Cre*^*+*^ and control *Cre*^-^ male (*Usp9x*^*fl/Y*^; *Tie2*^*Cre*^) mice. Generated pups were genotyped at P10 for *Cre*, and echocardiography was performed to assess cardiac function at 6 weeks, 16 weeks and 24 weeks. After the baseline echocardiography at 6 weeks, mice were fed a high fat western diet (Envigo# TD.88137) until 24 weeks of age and then euthanized to collect hearts for histological analyses. Transthoracic echocardiographic studies were performed after anesthetization with isoflurane (2% in 100% oxygen at a flow rate of 1.5 L/min) using a Vevo 2100 device equipped with 18–38 MHz linear-array transducer with a digital ultrasound system (Visualsonics Inc., Canada). Body temperature was monitored throughout the procedure using a rectal probe thermometer. Pulsed wave Doppler analysis of three consecutive cardiac cycles across the aortic valve was performed and averaged along parasternal long axis to obtain maximal transvalvular velocity. All echocardiographic experiments were performed by an experienced investigator blinded to the study design and animal genotypes.

After euthanasia, hearts were perfused with PBS, removed and fixed in 4% PFA at 4°C overnight. After fixation, the hearts were embedded in paraffin and sectioned. For gross histological analysis, tissue sections were stained with Russell-Movat Pentachrome stain (American MasterTech# KTRMP) to visualize collagen, elastin, muscle, mucin and fibrin and with Alizarin red (Acros Organics# 400480250) to visualize calcium deposition. For both staining, tissue sections were deparaffinized in xylene and rehydrated in grades of ethanol and in water. Russell-Movat Pentachrome staining was performed following the manufacturer’s protocol. For calcium deposition staining, tissue sections were incubated with 2% Alizarin-red in water for 5-7 minutes followed by washing in water. All images were visualized using the Olympus BX51 microscope attached with Olympus DP71 camera. Valve leaflet area was quantified using ImageJ software at least on three sections per aortic valve.

### Collection of aortic valve leaflets from human patients

Aortic valve leaflets were excised by the surgeons during aortic valve replacement surgery (primary surgical indication: calcific aortic stenosis) at Brigham and Women’s Hospital, Harvard Medical School, Boston, Massachusetts and processed as described before^57^. After excision, leaflets were placed in saline at 4°C for approximately 30 minutes and transported to the lab quickly (approximately within 15 minutes) in DMEM media (ThermoFisher# 10569044) without serum or antibiotics. Leaflets were then rinsed with cold PBS, and intact leaflets were placed into dry, 1.5 ml Eppendorf tubes, followed by a slow freezing in −80°C.

### Statistics

All experiment was performed at least in triplicate per treatment conditions. Data are presented as mean±SD. Two-tailed Student’s *t* test and Mann Whitney test were performed to determine statistical significance using the GraphPad Prism 8 software package. *P* value ≤ 0.05 were considered statistically significant.

### Study approval

This study was approved by the Institutional Animal Care and Use Committee at NCH (protocol AR16-00053) and conducted in accordance with the NIH’s *Guide for the Care and Use of Laboratory Animals* (National Academies Press, 2011). Human valve studies were approved by the Institutional Review Board (IRB) at Nationwide Children’s Hospital (IRB#07-00298), Brigham and Women’s Hospital (IRB#2011P001703)^21^. Informed consent was obtained from study subjects or parents of subjects less than 18 years of age. These aortic valves were received after surgery, without identification.

### Data availability

The authors declare that all supporting data are available within the article and its Supplemental Online Data. The plasmid constructs are commercially available from Addgene. The mouse models are commercially available from Jackson Laboratory, USA and Charles River Laboratories, Italy. Any other relevant data are available from the authors upon reasonable request. Source data are provided with this paper.

## Acknowledgements

The authors thank members of the Biomorphology Core and Adrianna P. Matos Nieves at NCH for histology support, and Dr. Marie Lockhart for assistance during preparation of pAVICs primary culture. The authors also thank Dr. Liwen Wang, Center for Proteomics and Bioinformatics, Case Western Reserve University for LC/MS-MS analysis.

## Author Contribution

VG and UM conceived the project and experiments were designed by UM and VG with input from BL, SEC, JL. UM performed and analyzed the experiments. The murine studies were performed by UM and EC. MB performed single cell RNA sequencing library preparation and siRNA mediated knockdown experiments. YU and MRM performed and analyzed the mouse echocardiograms. Bioinformatic analyses were performed by SNM. MCB and EA collected, graded and provided human tissues. SW generated and provided *Usp9x*^*loxp/loxp*^ mice. UM, VG, BL, SEC, JL interpreted the data. UM and VG wrote the manuscript with input from all authors.

## Additional Information

Online Figures 1-8

Online Table 1, 2

Online Video I, II

Online data I-IX

Uncropped blots

