## Supplemental file for "Nitric oxide prevents aortic valve calcification by S-nitrosylation of USP9X to activate NOTCH signaling"

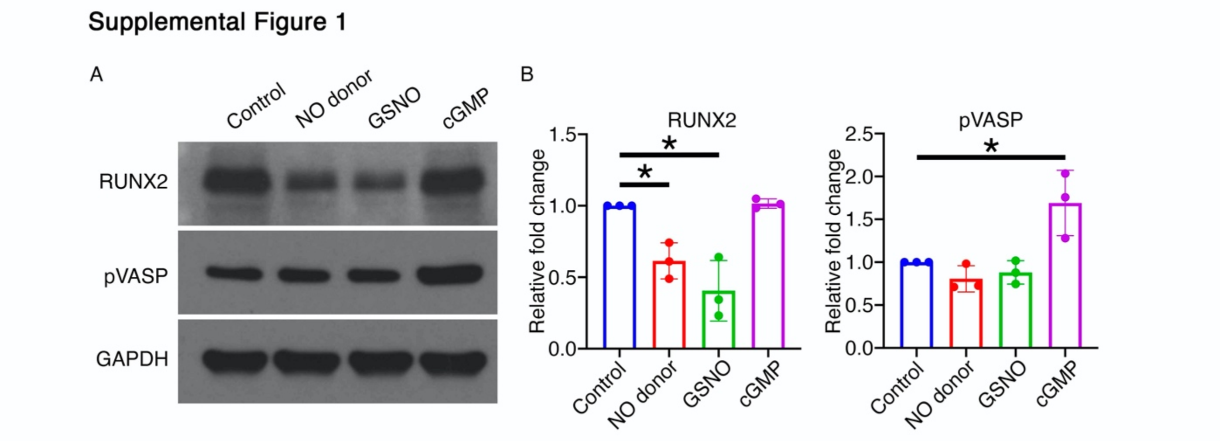


**Supplemental Figure 1.** (A) Immunoblot showing expression of RUNX2 and phosphorylated vasodilator-stimulated phosphoprotein (pVASP) in porcine aortic valve interstitial cells (AVICs) cultured for 5 days in the presence of detaNONOate (NO donor), GSNO (S-nitrosylating agent), 8-Br-PET-cGMP (cGMP analog) and compared to untreated cells (control). GAPDH served as loading control. Expression of RUNX2 protein was reduced with the addition of NO donor and GSNO, but not with 8-Br-PET-cGMP. Whereas pVASP expression was increased in presence of 8-Br-PET-cGMP but not with NO donor and GSNO (B) Graphs show quantifications after normalization to GAPDH. *, *P* value ≤ 0.05 (unpaired 2-tailed t-test).

Supplemental Figure 2

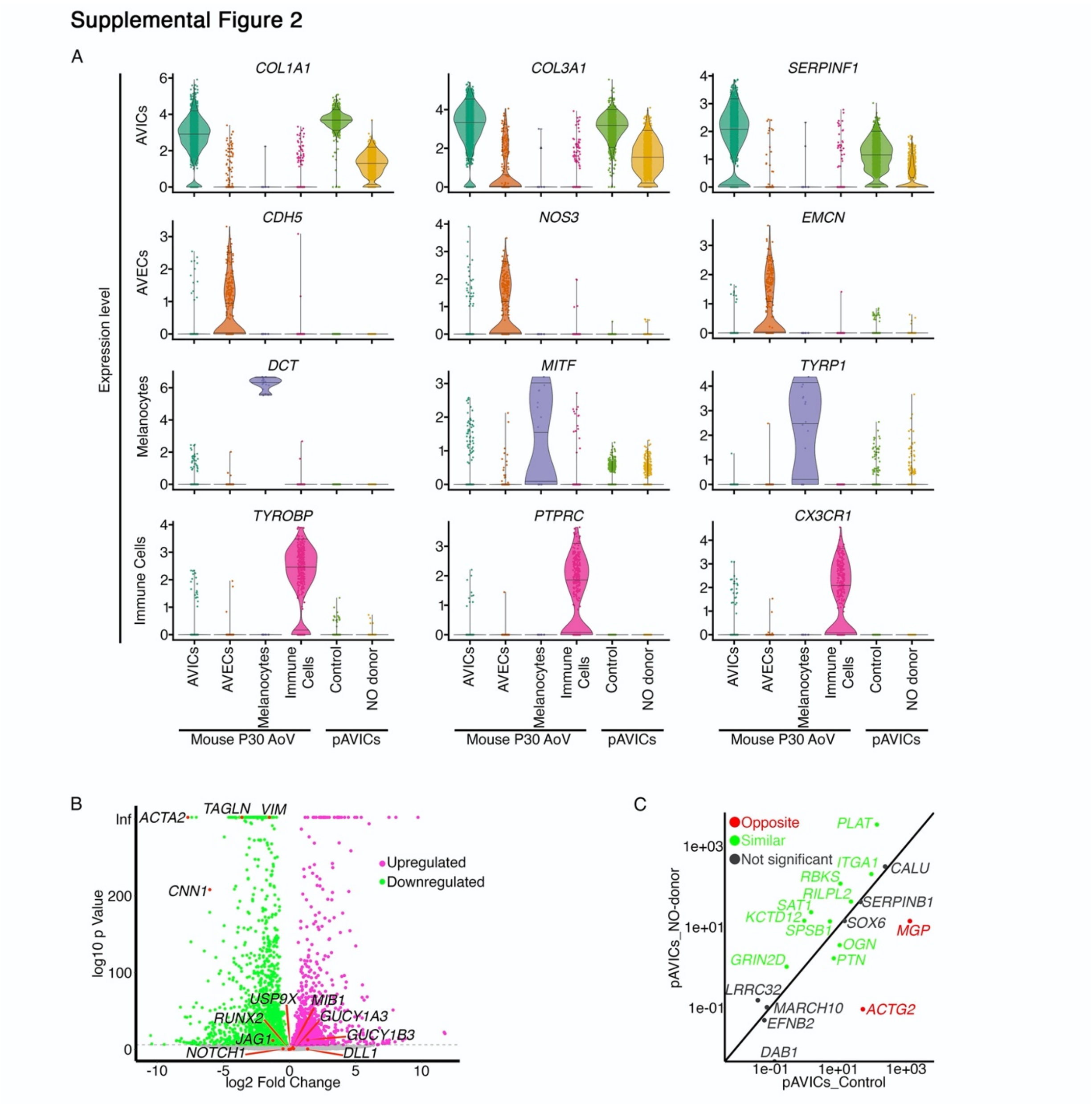


**Supplemental Figure 2.** Comparison of single cell RNA-seq (scRNA-seq) data from porcine aortic valve interstitial cells (AVICs) to previously published scRNA-seq data from one-month old mouse aortic valves[24]. (A) Violin plots show cultured porcine AVICs do not have substantial expression of genes that mark endothelial cells, melanocytes and immune cell but there is overlap with mouse AVICs. Horizontal lines in violin plot represent 0.05, 0.5 and 0.95 quantiles of expression of each gene. (B) Volcano plot of $\log_2$(Fold change) vs $-\log_{10}$(*P* value), showing differential expression of genes between NO donor treated and untreated cells. *ACTA2* is the most significantly downregulated gene after NO donor treatment. (C) Comparison of expression changes in cultured porcine AVICs treated with NO donor to previously identified NOTCH1 target genes[26]. Genes plotted above the identity line are upregulated and below the identity line are downregulated after NO donor exposure. Genes are color coded to indicate analogous changes (green) or opposite changes (red) between pAVICs and human AVECs. Genes that change in human aortic valve endothelial cells (AVECs) in response to sheer stress and Notch1 knockdown but have not changed in pAVICs are shown in grey. 10 out of the 16 genes detected in pAVICs show similar gene expression pattern.

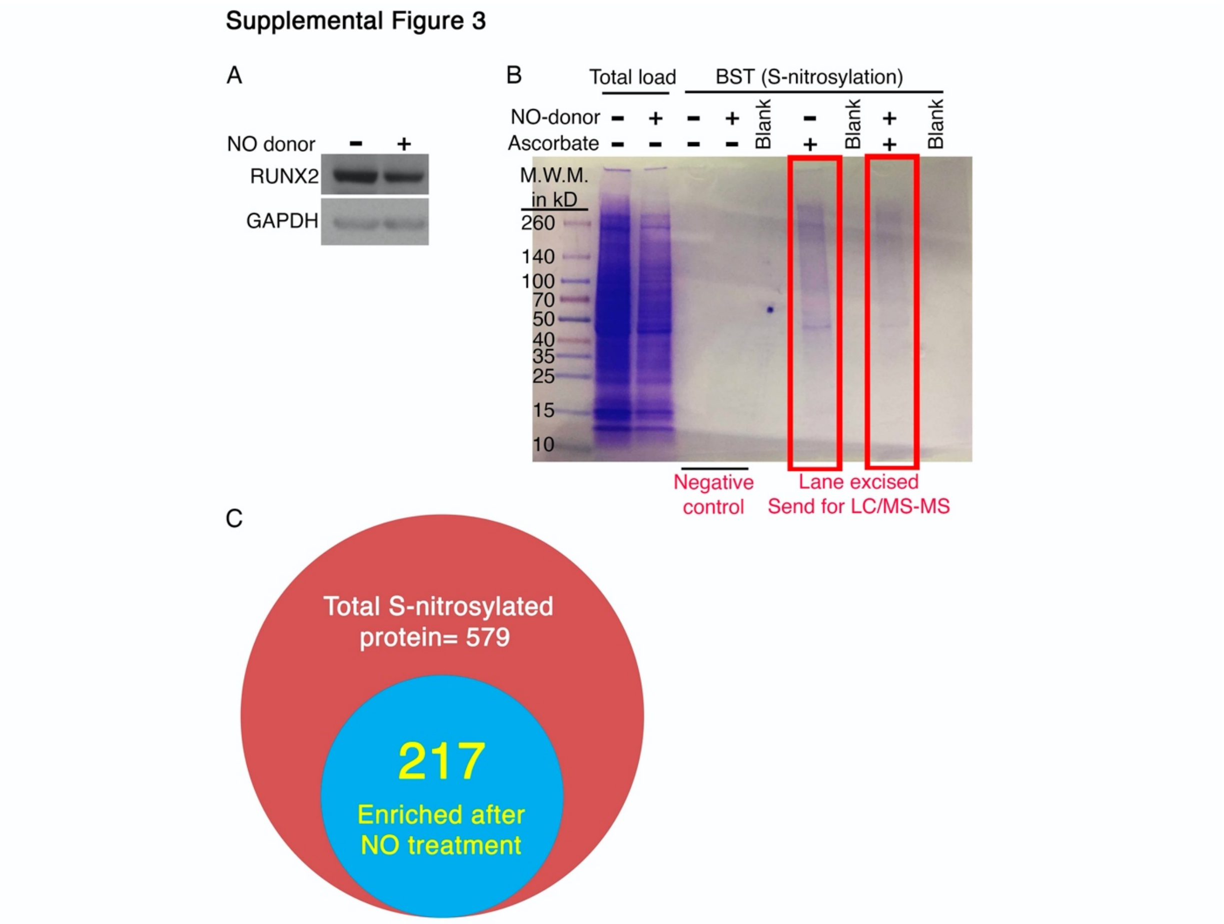


**Supplemental Figure 3.** S-nitrosylated proteins in rat aortic valve interstitial cells (AVICs) cultured in osteogenic media and exposed to NO donor were identified by using modified biotin switch technique (BST) followed by mass-spectrometry (MS). (A) Immunoblot demonstrates decreased expression of RUNX2 with addition of NO donor. GAPDH serves as loading control. (B) Representative SDS-PAGE stained with Imperial™ Protein Stain demonstrate bands of S-nitrosylated proteins in the presence of ascorbate. The no protein band in the absence of ascorbate serves as the negative control. Total load represents the protein extract that was used for the purification of S-nitrosylated proteins. Lanes of S-nitrosylated proteins were excised and subjected to LC/MS-MS analysis. (C)

Diagram representing the total number of identified S-nitrosylated proteins (579) number and those enriched after NO donor treatment (217).

Supplemental Figure 4

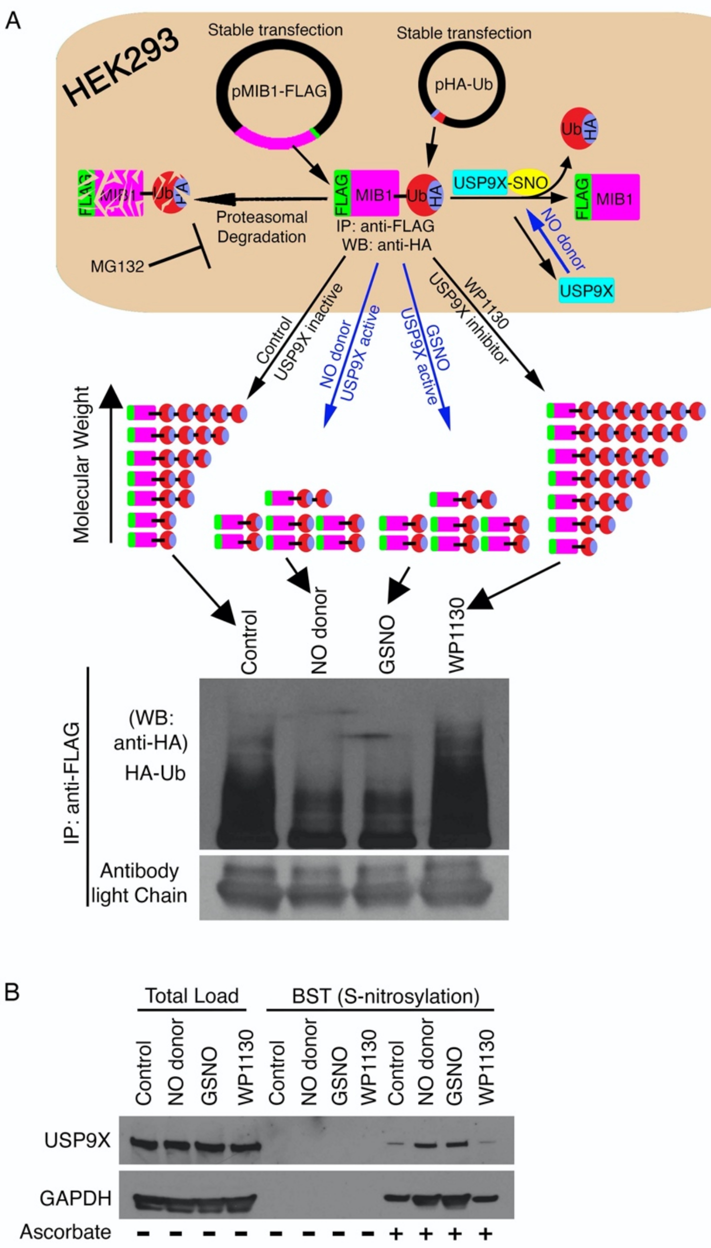


**Supplemental Figure 4.** (A) Schematic representation of the experimental design where HEK293 cells were engineered that stably expressed MIB1-FLAG and HA-Ub. In the presence of NO donor, S-nitrosylated USP9X deubiquitinates and stabilizes MIB1-FLAG. While in the absence of NO donor, MIB1-FLAG is degraded via the ubiquitin proteasome pathway due to the absence of S-nitrosylated USP9X. Immunoprecipitation (IP) with anti-

FLAG was performed after inhibiting proteasomal degradation with MG132. Immunoblot against HA-tag is shown. Diagram indicates the presence of various Ub-HA tags, according to the molecular weight under different culture conditions. Immunoblot demonstrates the predicted pattern with similar expression of MIB1-FLAG and GAPDH. (B) Modified biotin switch technique (BST) followed by immunoblot demonstrates S-nitrosylation of USP9X in HEK293 cells. No band in absence of ascorbate serves as the negative control. Total load represents the protein extract that was used for purification of S-nitrosylated proteins. GAPDH is a known S-nitrosylated protein and served as the positive control.

Supplemental Figure 5

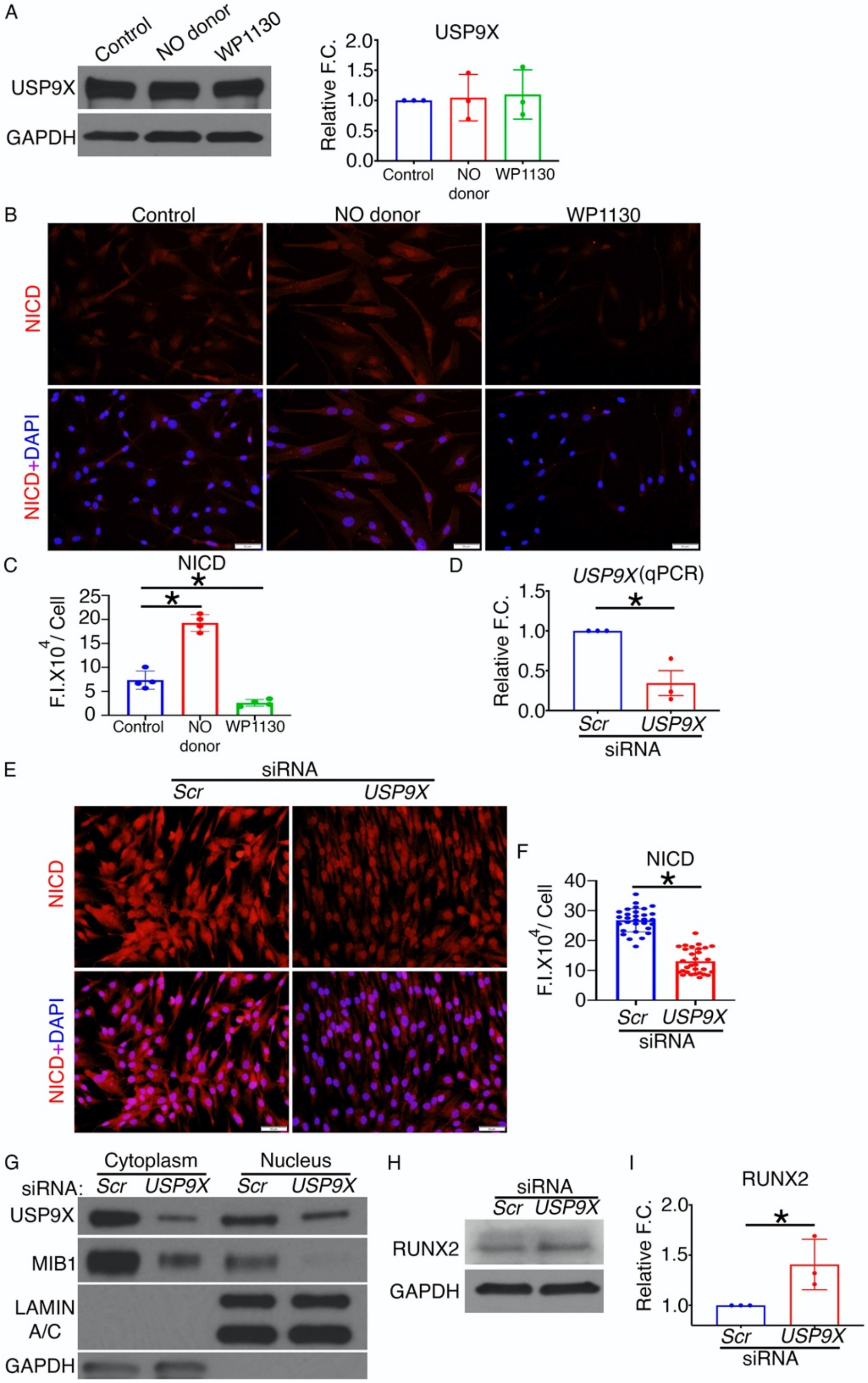


**Supplemental Figure 5.** Porcine aortic valve interstitial cells (AVICs) were cultured in the presence of detaNONOate (NO donor) or WP1130 (USP9X inhibitor) for 5 days and compared with an untreated control. (A) Expression of USP9X is shown by immunoblot in all culture conditions. GAPDH serves as a loading control. Graph shows quantification after normalization to GAPDH. (B) Immunofluorescence (IF) staining against NOTCH1 intracellular domain (NICD) is shown with counterstaining for nuclear DAPI porcine AVICs treated with NO donor and WP1130 in comparison to untreated cells (control). (C) Graph shows quantification of IF images normalized to nuclei. (D) Knockdown of *USP9X* with pooled *USP9X* specific siRNA in porcine (AVICs) compared to a scramble (*Scr*) control. RNA was isolated after 3 days of transfection and *USP9X* transcripts were significantly reduced by qPCR after *USP9X* knockdown compared to *Scr* (control). (E) IF staining against NICD after *USP9X* knockdown compared to *Scr* (control). Images are counterstained with nuclear DAPI. (F) Quantification of IF images demonstrate decreased NICD after *USP9X* knockdown. (G) In similar culture conditions, nuclear cytoplasmic fractionation followed by immunoblot shows reduced cytoplasmic and nuclear USP9X and MIB1 after *USP9X* knockdown. GAPDH (cytoplasmic) and LAMIN A/C (nuclear) serve as the loading controls. (H) Immunoblot demonstrates the expression of RUNX2 after *USP9X* knockdown. GAPDH serves as a loading control. (I) Quantification indicates significant increase of RUNX2 expression against GAPDH after *USP9X* knockdown. Scale bar: 50 µm. * represent p value ≤ 0.05 (2-tailed). For all IF, the Mann-Whitney test was performed and for WB and qPCR analyses, the unpaired two-tailed t-test were performed. F.I., fluorescence intensity; F.C., fold change.

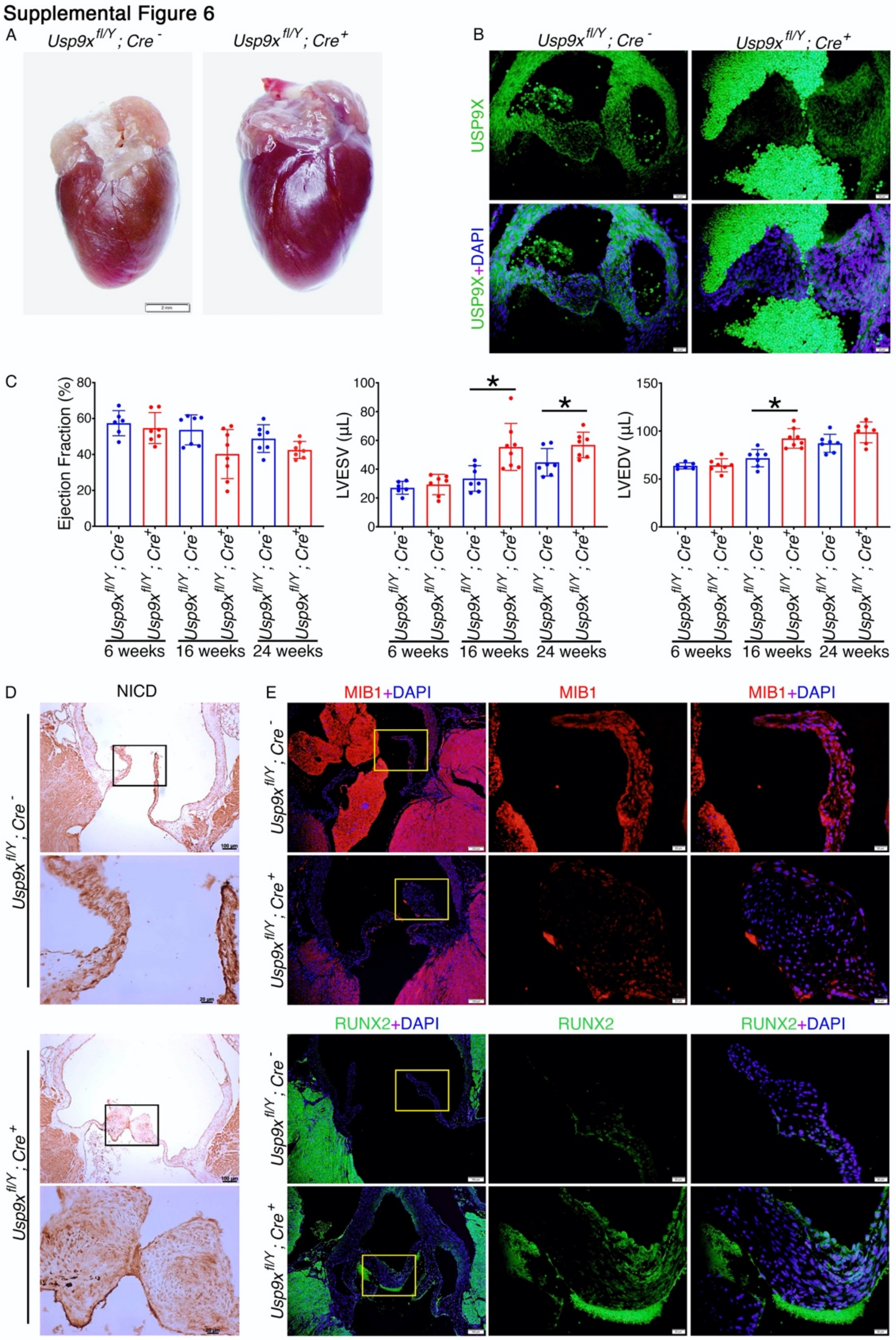


**Supplemental Figure 6.** (A) Representative image of hearts from 24 week old *Usp9x$^{fl/Y}$; Tie2$^{Cre+}$* and *Usp9x$^{fl/Y}$; Tie2$^{Cre-}$* are shown. (B) Immunofluorescence (IF) staining for Usp9x in aortic valve from E18.5 *Cre$^{+}$* male is shown in comparison to *Cre$^{-}$* male. (C) Ejection fraction, left ventricular end systolic volume (LVESV) and left ventricular end diastolic volume (LVEDV) from echocardiography are shown at 6, 16 and 24 weeks in *Cre*$^+$ males compared to *Cre*$^-$ males (controls). (D) Immunohistochemical (IHC) staining of NOTCH1 intracellular domain (NICD) demonstrates a decrease of NICD in 24 week old *Cre*$^+$ males compared to *Cre*$^-$ mice (control). Black rectangles are magnified at the bottom of each image. (E) IF demonstrates a decrease of MIB1 and an increase of RUNX2 in the aortic valve leaflets in 24-week-old *Cre*$^+$ males compared to *Cre*$^-$ males (control). Yellow rectangles are magnified at right of each image. Scale bar: (A) 2 mm; (B) 20 µm; (D and E) 100 µm and after magnification 20 µm. * represent p value $\leq$ 0.05 (2-tailed). For all experiments presented Mann-Whitney test was performed.

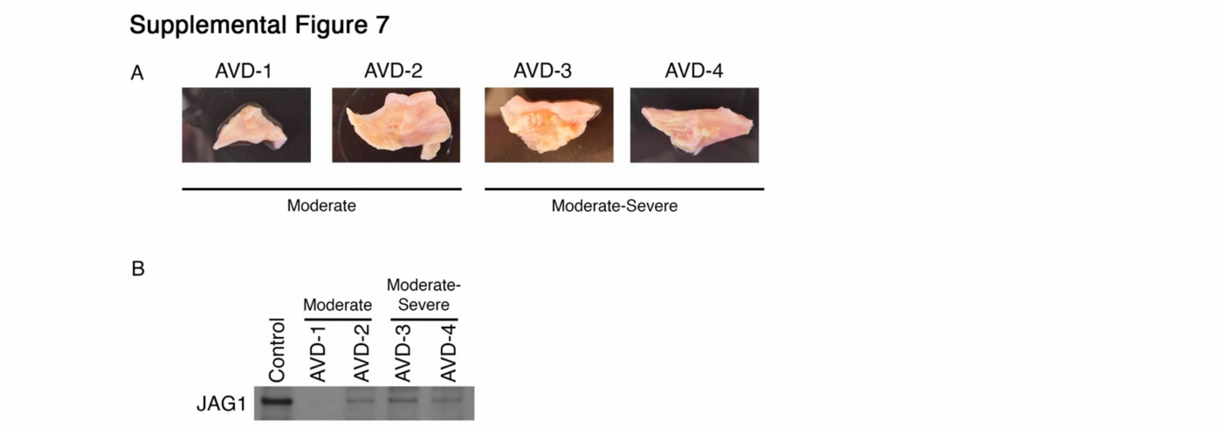


**Supplemental Figure 7.** (A) Explanted aortic valves (n=4) utilized in this study from adult patients categorized with moderate (n=2) and moderate-severe (n=2) calcification. (B) Immunoblot shows reduced expression of JAG1 in calcified aortic valves compared to non-calcified aortic valve from patient with aortic insufficiency.

Supplemental Figure 8

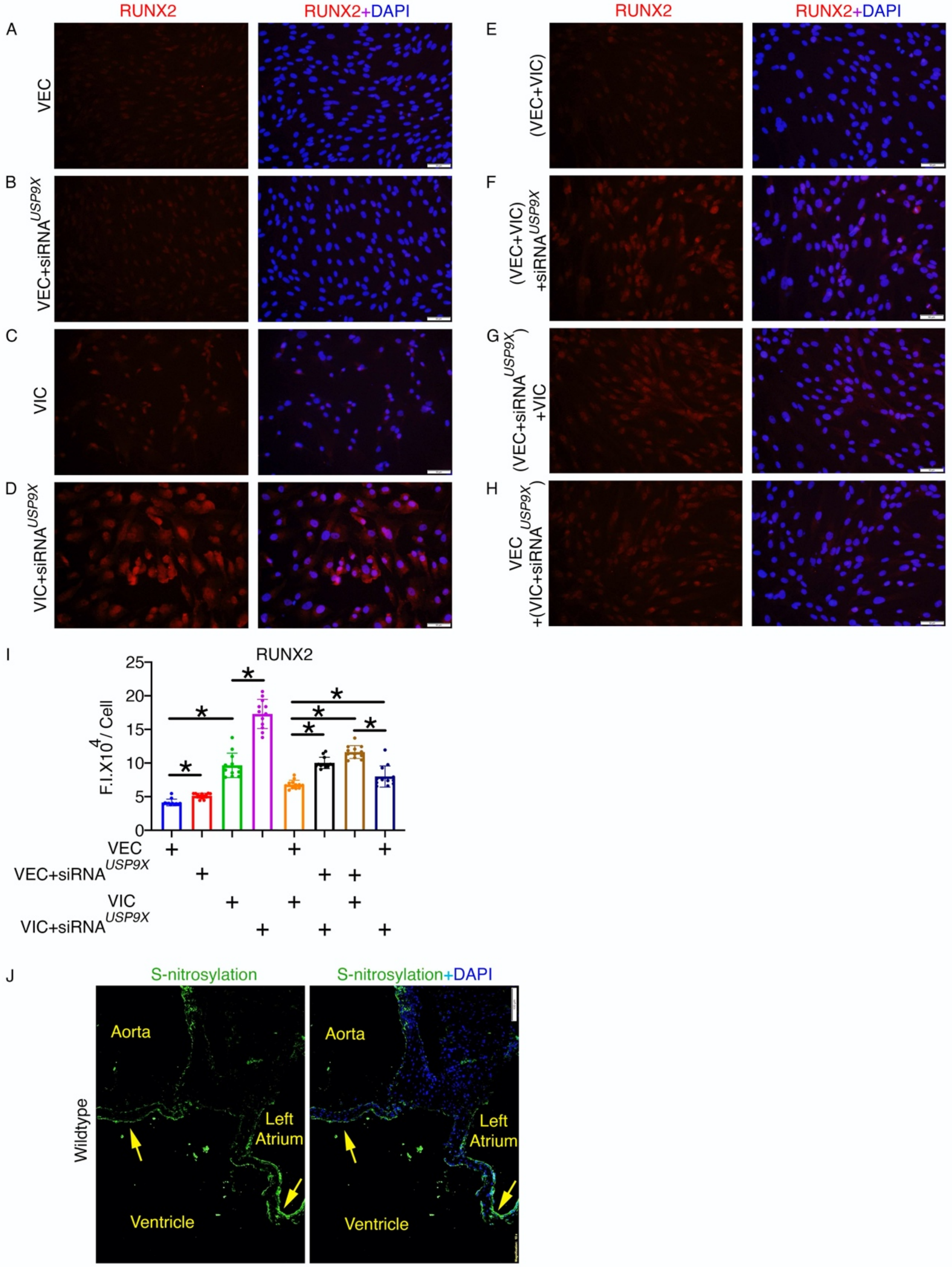


**Supplemental Figure 8.** Knockdown of *USP9X* by separate transfections of siRNA against *USP9X* in porcine aortic valve endothelial cells (AVECs) and interstitial cells

(AVICs). Twelve hours after transfection, cells were collected using TrypLE Express and co-cultured in the indicated combinations for 60 hours. (A-H) Immunofluorescent (IF) images demonstrate the expression of osteogenic marker, RUNX2, in untreated porcine AVICs or AVECs with *USP9X* knockdown as indicated. Images show nuclear DAPI counterstaining. Fluorescence intensity per cell was quantified. (I) Quantification of RUNX2 IF in different co-culture conditions, as indicated. (J) Modified biotin switch technique (BST) followed by IF was performed to visualize S-nitrosylated proteins in the adult mouse heart. Increased S-nitrosylation (arrow) was observed on the ventricular side of aortic valve leaflet and atrial side of mitral valve leaflet. * represent $P$ value ≤ 0.05 (2-tailed). For all experiments presented Mann-Whitney test was performed.

**Supplemental Table 1:**

**Primer Sequence:**

| Species Name | Gene Name | Primer Name | Sequence (5'>3') |
|---|---|---|---|
| *Sus scrofa* | *USP9X* | Sus_Usp9x_F1 | AGGCTCCTGATGGACAGTCT |
| | | Sus_Usp9x_R1 | TGTTCATCTGGGGGAGTTGC |
| | *GAPDH* | SusGAPDHqF | ACATGGCCTCCAAGGAGTAAGA |
| | | SusGAPDHqR | GATCGAGTTGGGGCTGTGACT |
| *Mus musculus* | *Cre* | Transgene F | GCGGTCTGGCAGTAAAAACTATC |
| | | Transgene R | GTGAAACAGCATTGCTGTCACTT |
| | | Internal+Control F | CTAGGCCACAGAATTGAAAGATCT |
| | | Internal+Control R | GTAGGTGGAAATTCTAGCATCATCC |

**Supplemental Table 2:**

**siRNA Sequence:**

| Species Name | Gene Name | Oligo Name | Sequence (5' >3') |
|---|---|---|---|
| *Sus scrofa* | *USP9X* | siRNA_susUsp9x_1 | AAGCACTTACTGAGTGGGAAT |
| | | siRNA_susUsp9x_2 | AAGGAATGGTATTCTTGCAAT |
| | | siRNA_susUsp9x_3 | AAGCAGGACAATGAGAGCAAT |
| | Scramble *(Scr)* | sus_scrSiRNA | AAGGTAAACGATTTATAGACT GCGATA |

**Online Video I:** Echocardiography demonstrates aortic flow in *Usp9x*$^{fl/Y}$*; Tie2*$^{Cre-}$ mouse without regurgitation.

**Online Video II:** Echocardiography demonstrates aortic flow in *Usp9x*$^{fl/Y}$*; Tie2*$^{Cre+}$ mouse with regurgitation.

**Online data I:** Average expression of top 100 differentially expressed genes in pAVICs after NO donor exposure compared to untreated control identified by scRNAseq.

**Online data II:** S-nitrosylated proteins identified by LC/MS-MS in control rat AVICS replicate 1.

**Online data III:** S-nitrosylated proteins identified by LC/MS-MS in control rat AVICS replicate 2

**Online data IV:** S-nitrosylated proteins identified by LC/MS-MS in control rat AVICS replicate 3

**Online data V:** S-nitrosylated proteins identified by LC/MS-MS in NO-treated rat AVICS replicate 1

**Online data VI:** S-nitrosylated proteins identified by LC/MS-MS in NO-treated rat AVICS replicate 2

**Online data VII:** S-nitrosylated proteins identified by LC/MS-MS in NO-treated rat AVICS replicate 3

**Online data VIII:** Total S-nitrosylated proteins in rat AVICS.

**Online data IX:** NO-enriched S-nitrosylated proteins in rat AVICs.
